# SPEAR: Spatial Epitranscriptomic Analysis by Mass Spectrometry Reveals Localized RNA Modification Remodeling During Glioma Progression

**DOI:** 10.64898/2026.10.13.758238

**Authors:** Stéphanie Mateus, Aurore Attina, Kévin Bigot, Valérie Rigau, Jérôme Vialaret, Christophe Hirtz, Amélie Darlix, Chris Planque, Alexandre David

## Abstract

RNA modifications shape cell identity and disease, yet their spatial organization within tissues has remained largely inaccessible to existing technologies. NGS-based methods can map a limited number of individual modifications with spatial resolution, while liquid chromatography-tandem mass spectrometry (LC-MS/MS) provides comprehensive and quantitative coverage of the epitranscriptome from bulk, homogenized tissue. No current approach combines histological resolution, broad modification coverage, quantitative accuracy, and the sensitivity required for microscale clinical specimens.

We present SPEAR (SPatial Epitranscriptomic Analysis by mass spectrometRy), a workflow coupling laser microdissection with a redesigned LC-MS/MS pipeline to enable quantitative epitranscriptomic profiling directly from formalin-fixed, paraffin-embedded (FFPE) tissue. This combination enables reproducible detection of dozens of RNA modifications from regions as small as 0.1 mm², establishing a spatial scale previously inaccessible to mass spectrometry-based approaches.

Applied to serial mouse brain sections, SPEAR resolved anatomically defined epitranscriptomic signatures across white and gray matter, establishing its spatial resolution and reproducibility in a controlled system. We then applied SPEAR to human diffuse gliomas containing, within individual specimens, both primary low-grade tumor regions and discrete foci undergoing transformation toward higher-grade malignancy. SPEAR resolved this transition directly, revealing a coordinated decrease in multiple methylated RNA modifications specifically within higher-grade regions, a spatially restricted signal that was masked by bulk analysis and reproducibly observed across patients despite underlying genetic heterogeneity. SPEAR establishes spatial epitranscriptomics as a tractable approach for molecular profiling of clinical tissue, providing a platform for biomarker discovery and molecular stratification based on the spatial organization of RNA modifications.

## INTRODUCTION

Chemical modifications of RNA, collectively referred to as the epitranscriptome, constitute a fundamental layer of post-transcriptional regulation. To date, more than 170 distinct RNA modifications have been identified across all RNA classes, in which they regulate RNA stability, splicing, translation, and other aspects of RNA metabolism, thereby shaping cell identity and phenotypic plasticity in health and disease (Jonkhout et al. 2017; Sordyl et al. 2026). Because many RNA modifications respond dynamically to environmental and cellular stresses, dysregulation of the epitranscriptome has increasingly been implicated in cancer evolution, cellular plasticity, and therapeutic resistance (Delaunay et Frye 2019). We previously demonstrated that LC-MS/MS-based quantitative profiling of RNA modifications generates tumor-state-associated molecular signatures with potential applications in cancer stratification and precision medicine (Relier et al. 2022). However, these analyses relied on bulk tissue measurements and therefore averaged signals across heterogeneous cellular populations, precluding the investigation of localized epitranscriptomic states associated with tissue architecture, microenvironmental context, and spatial clonal evolution (Venteicher et al. 2017; Darlix et al. 2020).

Although recent NGS-based approaches have enabled transcript-level mapping of individual RNA modifications, including m^6^A and inosine, with increasing spatial resolution (Xu et al. 2026), they are primarily designed to interrogate individual RNA marks rather than the epitranscriptome as a whole. Conversely, LC-MS/MS enables comprehensive and quantitative profiling of diverse RNA modifications but remains confined to bulk tissue analysis. A recent mass spectrometry imaging method can spatially map only free modified nucleosides in frozen tissue (Peng et al. 2026), which is distinct from profiling RNA-associated modifications and has limited clinical applicability, as patient biopsies are commonly preserved as formalin-fixed, paraffin-embedded tissue. Comprehensive spatial characterization of the epitranscriptome, especially for clinical applications, is therefore beyond the reach of current technologies.

Closing this gap requires substantial gains in analytical sensitivity, as laser-microdissected tissue regions contain orders of magnitude less RNA than conventional LC-MS/MS workflows typically require. While LC-MS/MS remains the gold standard for comprehensive, quantitative analysis of RNA modifications, its application to microscale samples is constrained by both chromatographic and instrumental limitations. Nucleoside analysis is particularly challenging given the high polarity of these compounds, which often results in poor reversed-phase retention. Many profiling methods therefore rely on buffered mobile phases to improve retention; however, non-volatile salts promote ion suppression and progressively contaminate the ion source, limiting sensitivity (Annesley 2003). Separately, analyte adsorption onto metallic LC surfaces can further compromise recovery of polar, low-abundance compounds when only trace material is available (Guimaraes et Bartlett 2023).

To address these constraints, we developed SPEAR (SPatial Epitranscriptomic Analysis by mass spectrometRy), an integrated workflow that combines laser microdissection of formalin-fixed, paraffin-embedded (FFPE) tissue sections with a tailored, ultrasensitive LC-MS/MS method, integrating a salt-free, acidic mobile phase, a polar-modified reversed-phase column, and an inert chromatographic system. SPEAR enables comprehensive and quantitative profiling of RNA modifications from microscale, histologically defined tissue regions while preserving spatial context. By delivering robust measurements of dozens of RNA marks from spatially resolved clinical specimens, SPEAR extends spatial epitranscriptomic profiling to clinically relevant tissue scales and opens new opportunities for the analysis of human clinical samples.

As a proof of concept, we first applied SPEAR to the mouse brain, where anatomically distinct gray- and white-matter regions provide a controlled model of spatial cellular heterogeneity. We then used it to study emerging high-grade foci in low-grade human glioma samples (Darlix et al. 2020). SPEAR demonstrated robustness and reproducibility in resolving tumor heterogeneity and mapped with high spatial precision the reorganization of the epitranscriptomic landscape during tumor progression. Altogether, this work provides the first spatial characterization of the human glioma epitranscriptome, demonstrates that RNA modifications distinguish regional malignant cell states within individual tumors, and establishes spatial epitranscriptomic profiling as a new technological platform for dissecting intratumoral evolution.

## DESIGN

Despite the growing recognition that RNA modifications regulate cell identity and cellular plasticity, their spatial organization within complex tissues remains largely inaccessible. Sequencing-based approaches provide transcript level information but typically target a limited subset of RNA modifications, whereas LC-MS/MS enables comprehensive and quantitative profiling but is conventionally performed on bulk tissue, sacrificing spatial information. SPEAR was designed to bridge this gap by combining laser microdissection (LMD) with ultrasensitive LC-MS/MS for quantitative epitranscroptomic profling of histologically defined FFPE tissue regions. To maximize sensitivity while preserving spatial information, we optimized RNA extraction from FFPE specimens, including proteinase K-mediated reversal of formaldehyde-induced crosslinks, and established the minimal tissue area compatible with robust and reproducible epitranscriptomic profiling. We first benchmarked SPEAR on serial mouse brain sections to assess spatial resolution and reproducibility across anatomically distinct regions. We then applied the workflow to archived human diffuse gliomas containing paired low- and higher-grade regions within the same surgical specimens, enabling within-patient comparisons while minimizing interpatient genetic variability. This design allowed us to demonstrate that SPEAR resolves tumor heterogeneity and reveals localized epitranscriptomic remodeling associated with focal malignant progression.

## RESULTS

### Development of a salt-free chromatographic method for spatial epitranscriptomics

Because laser-microdissected FFPE samples yield only minute amounts of RNA, increased LC-MS/MS sensitivity was required to enable spatially resolved epitranscriptomic profiling. We therefore developed a salt-free chromatographic workflow using a mobile phase containing 0.5% acetic acid in combination with a Synergi Hydro-RP stationary phase. This configuration was selected to maintain the retention and separation of polar nucleosides in the absence of ammonium acetate and was benchmarked against our previously established ammonium acetate-based method (Amalric et al. 2024). Despite the complete removal of ammonium acetate from the mobile phase, the optimized method maintained overall chromatographic performance comparable to that of the reference conditions (**Figure 1A**). Most nucleosides displayed only moderate shifts in retention time, although the extent of these changes varied across nucleoside families. Guanosine- and uridine-derived modifications exhibited particularly stable chromatographic behavior between the two methods, whereas adenosine- and cytidine-derived modifications showed more pronounced shifts in retention time (**Figure 1B**). Importantly, the salt-free workflow substantially increased MS signal intensities for most nucleosides. Signal intensities increased by approximately 4-, 8-, 11-, and 8-fold for adenosine, cytidine, guanosine, and uridine-derived nucleosides, respectively (**Figure 1C**). The improvement was particularly pronounced for uridine-derived modifications, with signal enhancements reaching 12-fold for m³U and approximately 11-fold for both Um and ncm⁵U (**Figure 1D**). Conversely, a limited subset of adenosine-derived (m¹A and i⁶A) and guanosine-derived (m⁷G and m², ⁷G) modifications showed higher signal intensities under the ammonium acetate-based conditions, indicating that the impact of chromatographic composition on MS response is dependent, at least in part, on nucleoside structure (**Supplementary Figure 1**). Transfer of the optimized method to a bioinert LC platform further enhanced MS signal intensities for nearly all modified nucleosides, with signal gains of at least two-fold for most analytes, except ho⁵U. This effect was particularly pronounced for low-abundance sulfur-containing modifications, for which signal intensities increased by 3.4-fold for mcm⁵s²U and 7.8-fold for s²C **(Figure 1E)**. These improvements are consistent with reduced non-specific adsorption of nucleosides to metallic surfaces, thereby potentially increasing analyte recovery prior to ionization and detection.

**Figure 1:**
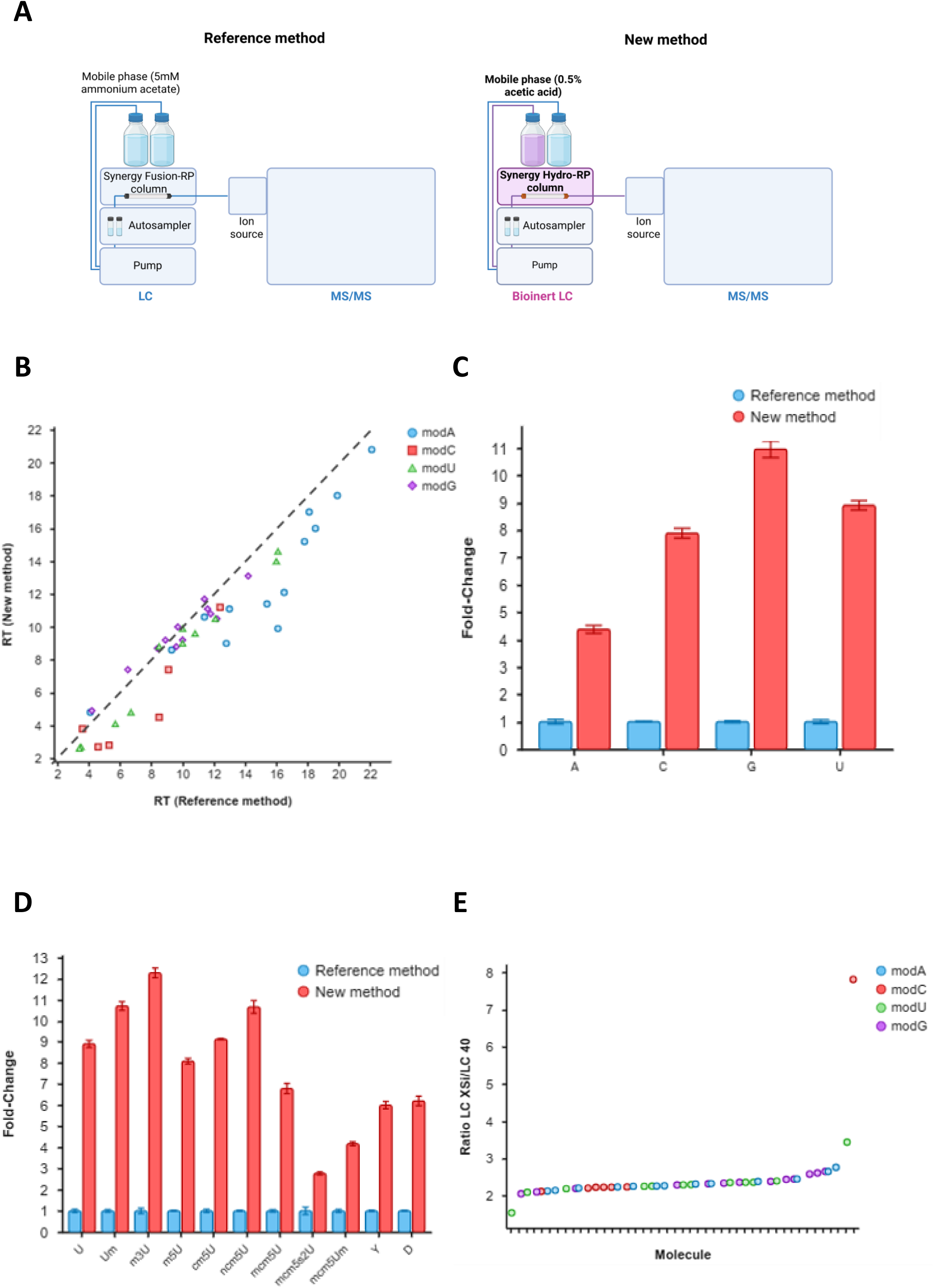
Optimization of chromatographic conditions for salt-free nucleoside analysis. A. Development of an optimized bioinert LC-MS/MS workflow for modified nucleoside analysis. Comparison of the reference (left) and optimized (right) workflows, highlighting the replacement of the conventional LC column and mobile phase with a Synergy Hydro-RP bioinert column and optimized mobile-phase composition. Common analytical steps include autosampling, chromatographic separation, ionization, and MS/MS detection. B. Comparison of nucleoside retention times obtained using the reference method (5 mM ammonium acetate, Synergi Fusion-RP column) and the new method (0.5% acetic acid, Synergi Hydro-RP column). The dashed line represents the line of identity (y = x). C. Comparison of fold-change values obtained using the new method relative to the reference method for canonical nucleosides. Fold-change values were calculated relative to the reference method, which was set to 1. Data are presented as mean ± SD. D. Comparison of fold-change values obtained using the new method relative to the reference method for uridine and selected uridine-derived modifications. Fold-change values were calculated relative to the reference method, which was set to 1. Bars represent mean values, and error bars indicate SD. E. Effect of transferring the method to a bioinert LC system on nucleoside detection sensitivity. Data are presented as the ratio of peak areas obtained with the bioinert LC system (LC XSi) to those obtained with the conventional LC-40 system.

Collectively, the combination of a 0.5% acetic acid mobile phase, a Synergi Hydro-RP stationary phase, and a bioinert LC system provided an optimized analytical platform that preserved robust chromatographic performance while substantially enhancing MS sensitivity. This workflow therefore enabled the sensitive detection of modified nucleosides from the minute RNA amounts obtained by laser microdissection of FFPE tissue, providing a suitable analytical foundation for spatially resolved epitranscriptomic profiling.

### Proteinase K-assisted RNA extraction enables sensitive SPEAR profiling from FFPE tissues

Formalin-fixed, paraffin-embedded (FFPE) tissue constitutes the standard preservation method for clinical biopsy specimens, enabling long-term storage at room temperature and providing an extensive source of archived material available for retrospective studies and biomarker discovery. Because SPEAR was designed with future clinical applications in mind, establishing an efficient RNA extraction procedure from FFPE tissue was an essential prerequisite. However, RNA extraction from FFPE samples remains technically challenging, as formaldehyde-induced cross-linking and RNA fragmentation can substantially impair nucleic acid recovery and compromise downstream molecular analyses (Kashofer et al. 2013). To establish the RNA extraction workflow for SPEAR, we systematically compared three RNA extraction strategies using FFPE mouse brain sections: a conventional TRIzol-based protocol (Protocol 1), a modified TRIzol protocol incorporating proteinase K digestion before RNA extraction (Protocol 2), and a silica column-based protocol including the same proteinase K digestion step (Protocol 3) (**Figure 2A**).

**Figure 2:**
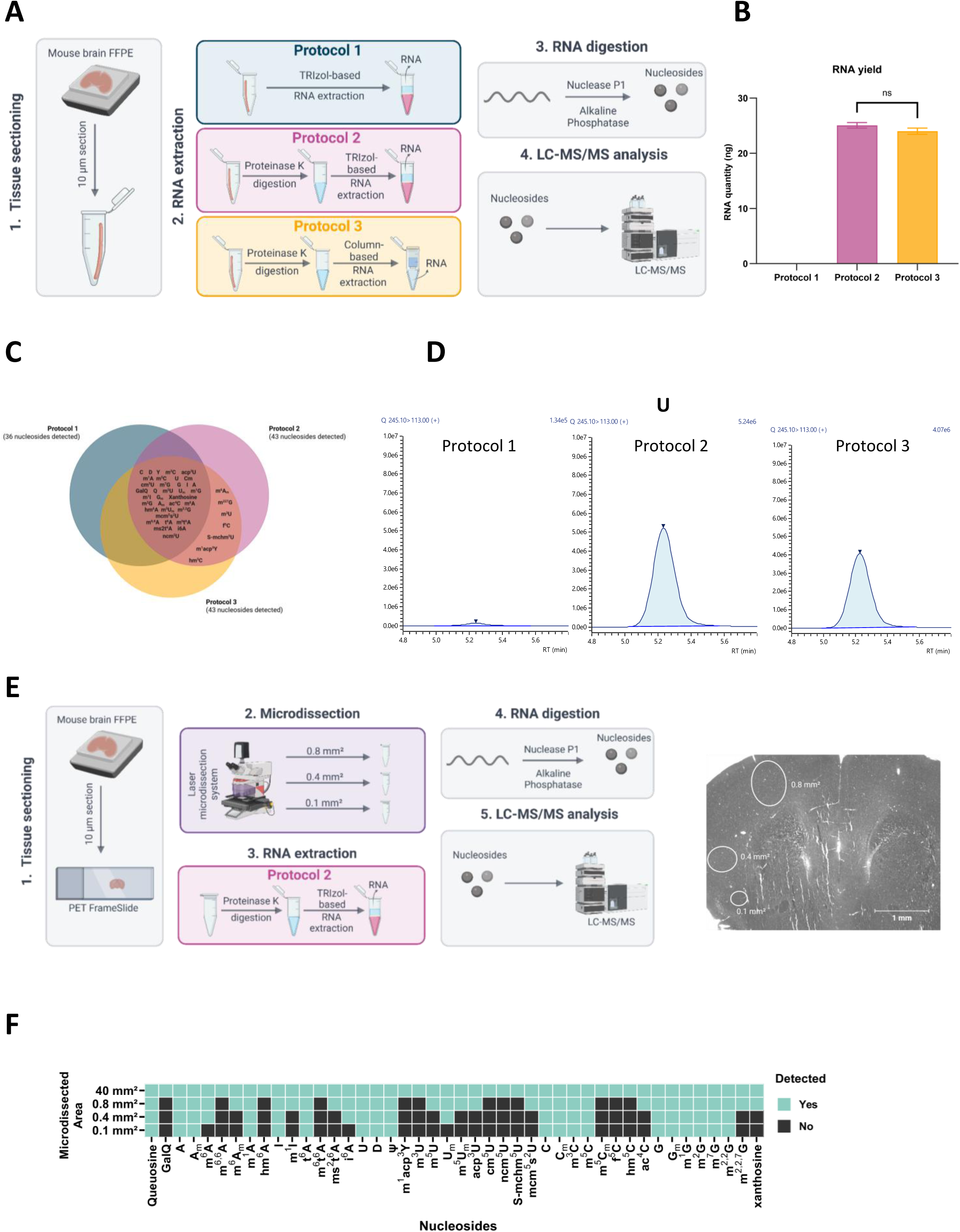
Optimization of the SPEAR workflow for RNA extraction and LC-MS/MS analysis from FFPE tissues. A. Schematic overview of the workflow: FFPE tissue sections were subjected to (1) sectioning, (2) RNA extraction using three different protocols: TRIzol extraction alone (protocol 1); proteinase K digestion followed by TRIzol extraction (protocol 2); or proteinase K digestion followed by silica column purification (protocol 3), (3) enzymatic digestion of RNA into individual nucleosides and (4) quantitative LC-MS/MS analysis. B. RNA yield obtained using the three extraction protocols, as measured using the Qubit fluorometric assay. Data are presented as mean ± SD from three independent experiments. Statistical significance was assessed using the Mann-Whitney U test (ns, p>0.05). C. Venn diagram showing the overlap of RNA modifications detected by LC-MS/MS across the three extraction protocols. D. Representative extracted ion chromatograms illustrating the uridine (U) signal intensity obtained with the different RNA extraction protocols. E. Schematic overview of the SPEAR workflow: FFPE tissues were subjected to (1) sectioning, (2) laser microdissection (LMD), (3) RNA extraction using Protocol 2, (4) enzymatic digestion of RNA into individual nucleosides and (5) quantitative LC-MS/MS analysis. F. Heatmap summarizing the detection of individual RNA modifications in whole FFPE tissue sections and in progressively smaller laser-microdissected areas, illustrating the analytical sensitivity of the SPEAR workflow.

Proteinase K digestion markedly improved RNA recovery from FFPE tissue. Whereas Protocol 1 yielded no detectable RNA, Protocols 2 and 3 recovered 25.07 ± 0.50 ng and 24.00 ± 0.57 ng of total RNA, respectively, from an entire FFPE mouse brain section (approximately 40 mm²) (**Figure 2B**). RNA quality was subsequently assessed by RT-qPCR using 28S rRNA, *Actb*, and *Gapdh*, with expression levels normalized to 18S rRNA. Consistent with the RNA recovery measurements, Protocol 1 failed to support reliable amplification, whereas Protocols 2 and 3 produced highly comparable expression profiles for all reference transcripts (**Supplementary Figure 2A**).

We next assessed whether the extraction procedure affected the analytical performance of SPEAR (**Supplementary Table 1**). Conventional extraction using Protocol 1 enabled the detection of 36 nucleosides, whereas incorporation of proteinase K digestion increased the number of detected nucleosides to 43, corresponding to the detection of seven additional nucleosides with both Protocols 2 and 3 (**Figure 2C**). Moreover, Protocol 2 consistently produced higher MS signal intensities than either Protocol 1 or Protocol 3 (**Figure 2D, Supplementary Figure 2B**), indicating improved recovery and/or preservation of modified nucleosides during extraction. Together, these results identify proteinase K digestion as a critical step in the SPEAR workflow, substantially increasing the number of detectable RNA modifications and improving analytical sensitivity in FFPE-derived samples. By facilitating the release of RNA from formaldehyde-cross-linked tissue, proteinase K treatment may alleviate a major limitation associated with FFPE-derived material. Because Protocol 2 achieved analytical performance comparable to that of Protocol 3 while avoiding the additional cost and handling associated with silica-column purification, it was selected as the standard RNA extraction procedure for all subsequent SPEAR analyses.

### Defining the analytical sensitivity of SPEAR across decreasing microdissected tissue areas

We next sought to define the minimal microdissected tissue area compatible with robust quantitative epitranscriptomic profiling using SPEAR. Starting from an entire FFPE mouse brain section of approximately 40 mm² (10 µm thickness), from which 44 nucleosides were quantified, we progressively reduced the laser-microdissected area to 0.05 mm² (**Figure 2E**). Progressive reduction of the microdissected tissue area produced a corresponding decrease in nucleoside coverage, with uridine-derived RNA modifications exhibiting the greatest loss of detectability, consistent with the inherently lower detectability of this class of nucleosides (Heiss et al. 2021).

Despite the progressive reduction in microdissected tissue area, SPEAR maintained quantitative analytical performance across sub-millimeter histological regions (**Figure 2F**). Microdissection of 0.8 mm² enabled detection of 32 nucleosides (73% of the full-section profile), whereas 0.4 mm² still enabled quantification of 22 nucleosides (50%). Even at 0.1 mm², corresponding to the detection of 19 nucleosides (43% of the full-section profile), SPEAR generated reproducible quantitative epitranscriptomic profiles, although with reduced nucleoside coverage compared with larger tissue areas. Only when the microdissected tissue area was reduced to 0.05 mm² did most nucleoside signals fall below the detection threshold.

Together, these findings establish the practical lower analytical range of SPEAR and demonstrate that multiplexed, quantitative epitranscriptomic profiling can be achieved from sub-millimeter histological regions. This level of analytical sensitivity enables the interrogation of highly limited tissue regions and makes SPEAR particularly well suited for laser-microdissected FFPE samples and other limited clinical specimens.

### Benchmarking the spatial resolving power of SPEAR using anatomically defined brain regions

To benchmark the spatial resolving capacity of SPEAR, we applied the workflow to FFPE mouse brain tissue, where well-defined anatomical compartments provide a controlled model of regional molecular heterogeneity. We established a serial-section strategy in which one section was stained and annotated to define histologically distinct regions of interest (ROIs), while the adjacent unstained section was subjected to laser microdissection followed by quantitative LC-MS/MS analysis (**Figure 3A**).

**Figure 3:**
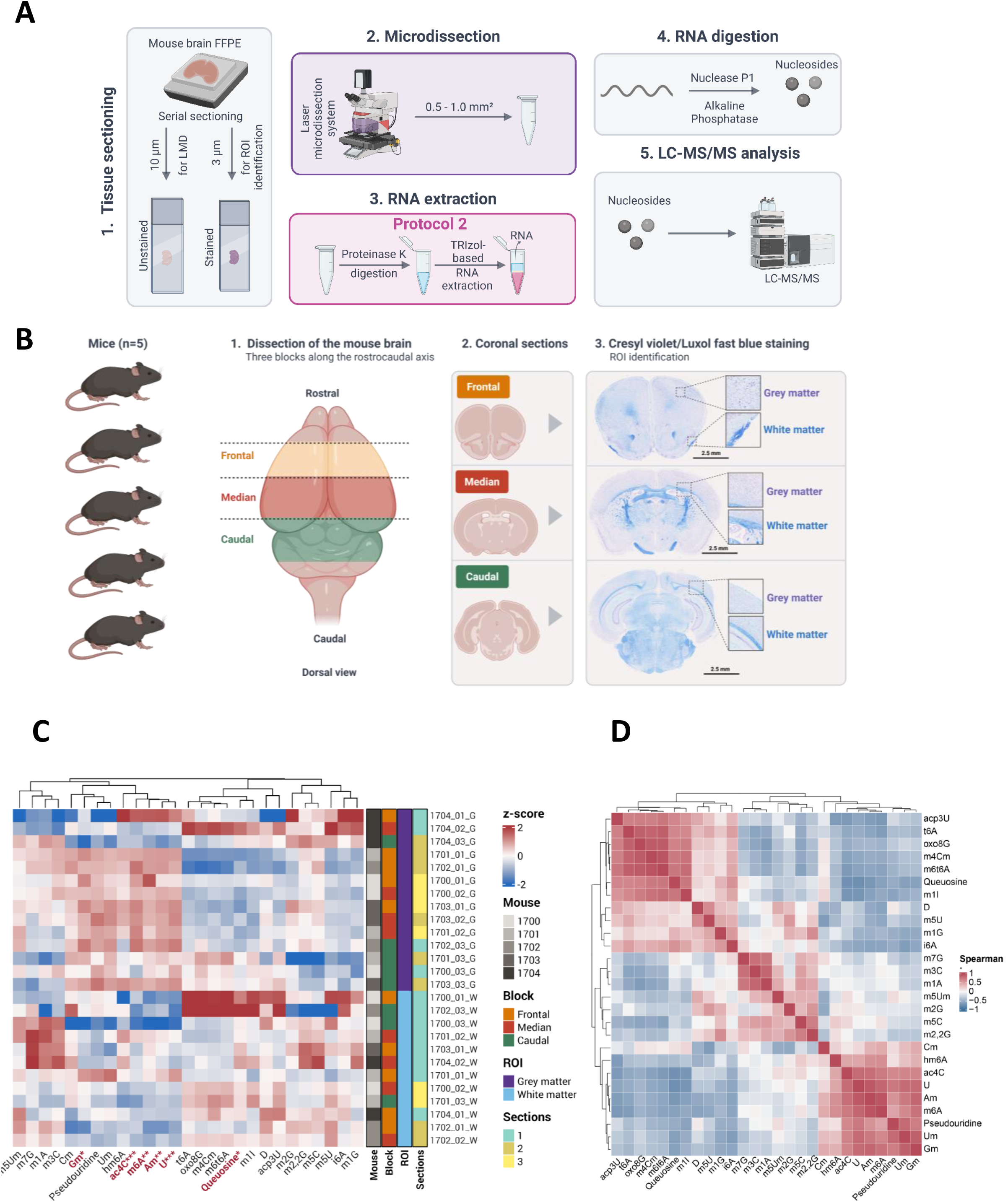
Spatial epitranscriptomic profiling of anatomically distinct regions of the mouse brain using SPEAR. A. Schematic overview of the SPEAR workflow: (1) regions of interest (ROIs) were identified in FFPE mouse brain sections, (2) isolated by laser microdissection (LMD), (3) processed for RNA extraction using Protocol 2, (4) enzymatically digested to individual nucleosides, and (5) analyzed by quantitative LC-MS/MS. B. Anatomical localization of the ROIs in coronal mouse brain sections and representative Luxol Fast Blue/Cresyl Violet-stained sections. White matter is stained blue, whereas gray matter is stained purple. C. Heatmap showing relative RNA modification abundance across samples collected from gray and white matter. RNA modification abundance is presented as the mean of one to three individual sections, when available. Modifications identified as ROI-associated by linear modeling are highlighted in red. Samples were ordered by ROI, and modifications were ordered according to the significance of the zone effect. Hierarchical clustering was performed within each predefined group using Euclidean distance and complete linkage. D. Heatmap showing Spearman correlations of RNA modification abundance across sections collected from gray and white matter.

The workflow was evaluated across five adult mouse brains, each represented by three independent FFPE tissue blocks. From each block, three consecutive coronal sections were analyzed, providing technical triplicates across serial sections. Within each section, anatomically defined white matter and gray matter regions were independently microdissected (**Figure 3B**). Mixed white/gray matter interface regions were additionally collected for exploratory analyses (**Supplementary Figure 3A**). For each ROI, laser capture was optimized to recover 0.5-1.0 mm² of tissue, either through a single dissection or through the pooling of multiple smaller microdissected areas into a single collection tube. Overall, this experimental design generated 90 spatially resolved samples from white and gray matter regions, and 135 samples when mixed white/gray matter interface regions were included. The white and gray matter samples constituted the primary dataset for comparative statistical analyses, whereas the mixed interface regions were analyzed separately as an exploratory dataset.

White and gray matter were selected as benchmark regions because they represent anatomically, functionally, and cellularly distinct brain compartments, enriched respectively in myelinated axonal tracts and glial cells versus neuronal cell bodies and synaptic structures (Maldonado et Alsayouri 2023). The mixed white/gray matter interface regions were analyzed separately to explore spatial transitions between these anatomically distinct compartments (**Supplementary Figure 3A**).

Raw LC-MS/MS data were processed using a dedicated analytical pipeline incorporating signal-quality filtering, background correction, missing value handling, probabilistic quotient normalization (PQN), and multivariate statistical analysis (STAR Methods). Samples displaying insufficient nucleoside coverage or globally reduced analytical signal were excluded according to predefined quality-control criteria, generating a high-confidence dataset for spatial epitranscriptomic analyses (**Supplementary Figure 3B, Supplementary Table 2**).

Principal component analysis separated most gray- and white-matter samples along PC2 (**Supplementary Figure 4**), and heatmap visualization revealed anatomical compartment-associated patterns of RNA-modification abundance (**Figure 3C and Figure 3D**). Consistent with these observations, linear models incorporating block, mouse, and anatomical compartment as fixed effects identified 12 nucleosides that were significantly associated with anatomical compartment, while no significant effect of block was detected across the epitranscriptome. Among the compartment-associated nucleosides, queuosine was more abundant in white matter, whereas ac^4^C, U, A_m_, m^6^A, and G_m_ were more abundant in gray matter (**Figure 3C**, **Supplementary Table 3**). The exploratory analysis of mixed white/gray matter interface regions showed intermediate patterns for several compartment-associated modifications, consistent with a gradual transition in epitranscriptomic profiles across tissue boundaries (**Supplementary Figure 3A**).

Together, these results demonstrate that SPEAR captures biologically meaningful regional differences in the epitranscriptomic landscape while maintaining robustness across independent animals, tissue blocks, and serial FFPE sections. This validation establishes the analytical foundation for applying SPEAR to spatially heterogeneous human pathological specimens.

### SPEAR captures intratumoral heterogeneity in human diffuse gliomas

Having established the analytical performance of SPEAR in FFPE mouse brain tissue, we next evaluated whether the workflow could resolve clinically relevant spatial epitranscriptomic heterogeneity in archived human diffuse gliomas, where focal malignant transformation provides a unique model of intratumoral evolution. We assembled a retrospective cohort of FFPE astrocytomas and oligodendrogliomas resected between 2012 and 2016, each containing one or more histologically confirmed grade 2+ or WHO grade 3/4 (malignant transformation) foci embedded within an otherwise WHO grade 2 tumor (**Table 1**). Grade 2+ foci are defined as foci exhibiting high cellular density, high vascular density, or minimal endothelial proliferation, while not meeting the criteria for grade 3 or grade 4 (Pedeutour-Braccini et al. 2015). Foci were identified on tumor sections through expert neuropathological review (V. R.), who annotated ROIs corresponding both to the main WHO grade 2 tumor and to each higher grade (2+, 3 or 4) transformation focus. These ROIs were independently annotated on hematoxylin and eosin (H&E)-stained sections, and subsequently isolated from adjacent unstained sections by laser microdissection and analyzed using SPEAR (**Figure 4A**).

**Table 1:**
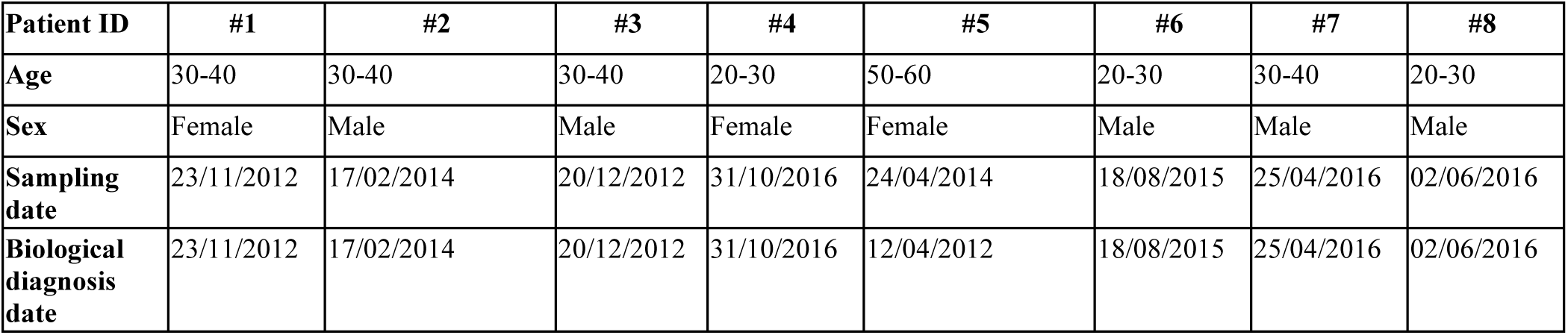
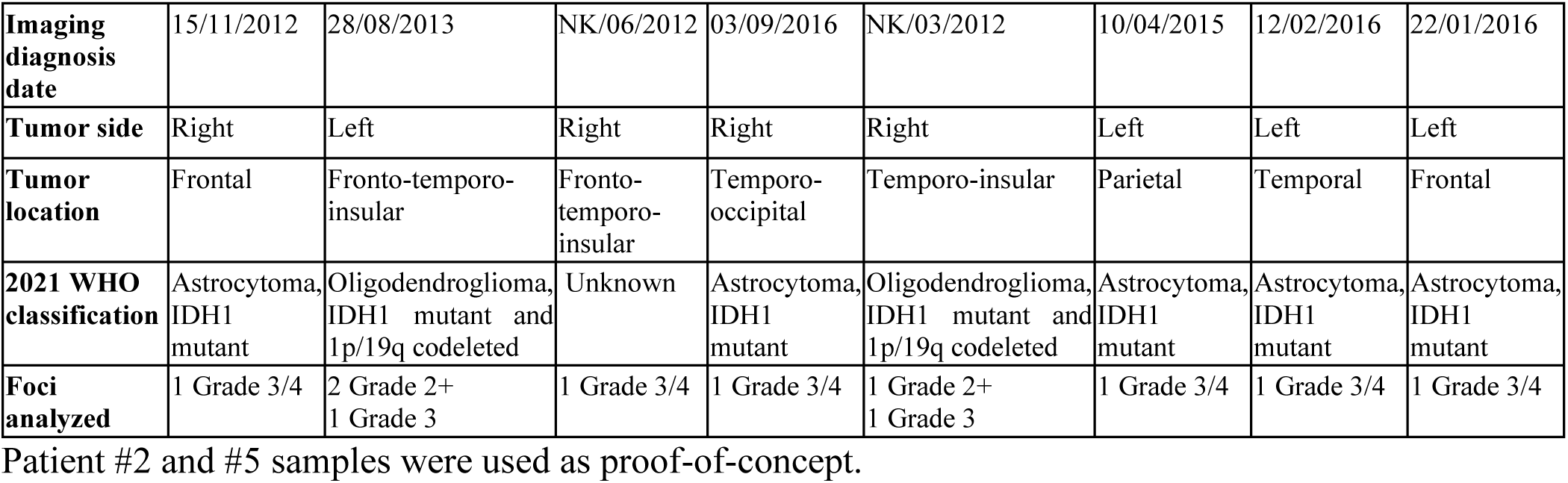
Clinical and pathological characteristics of the glioma cohort.

| Patient ID | #1 | #2 | #3 | #4 | #5 | #6 | #7 | #8 |
| --- | --- | --- | --- | --- | --- | --- | --- | --- |
| Age | 30-40 | 30-40 | 30-40 | 20-30 | 50-60 | 20-30 | 30-40 | 20-30 |
| Sex | Female | Male | Male | Female | Female | Male | Male | Male |
| Sampling date | 23/11/2012 | 17/02/2014 | 20/12/2012 | 31/10/2016 | 24/04/2014 | 18/08/2015 | 25/04/2016 | 02/06/2016 |
| Biological diagnosis date | 23/11/2012 | 17/02/2014 | 20/12/2012 | 31/10/2016 | 12/04/2012 | 18/08/2015 | 25/04/2016 | 02/06/2016 |
| **Imaging diagnosis date** | 15/11/2012 | 28/08/2013 | NK/06/2012 | 03/09/2016 | NK/03/2012 | 10/04/2015 | 12/02/2016 | 22/01/2016 |
| **Tumor side** | Right | Left | Right | Right | Right | Left | Left | Left |
| **Tumor location** | Frontal | Fronto-temporo-insular | Fronto-temporo-insular | Temporo-occipital | Temporo-insular | Parietal | Temporal | Frontal |
| **2021 WHO classification** | Astrocytoma, IDH1 mutant | Oligodendroglioma, IDH1 mutant and 1p/19q codeleted | Unknown | Astrocytoma, IDH1 mutant | Oligodendroglioma, IDH1 mutant and 1p/19q codeleted | Astrocytoma, IDH1 mutant | Astrocytoma, IDH1 mutant | Astrocytoma, IDH1 mutant |
| **Foci analyzed** | 1 Grade 3/4 | 2 Grade 2+<br>1 Grade 3 | 1 Grade 3/4 | 1 Grade 3/4 | 1 Grade 2+<br>1 Grade 3 | 1 Grade 3/4 | 1 Grade 3/4 | 1 Grade 3/4 |

Patient #2 and #5 samples were used as proof-of-concept.

**Figure 4:**
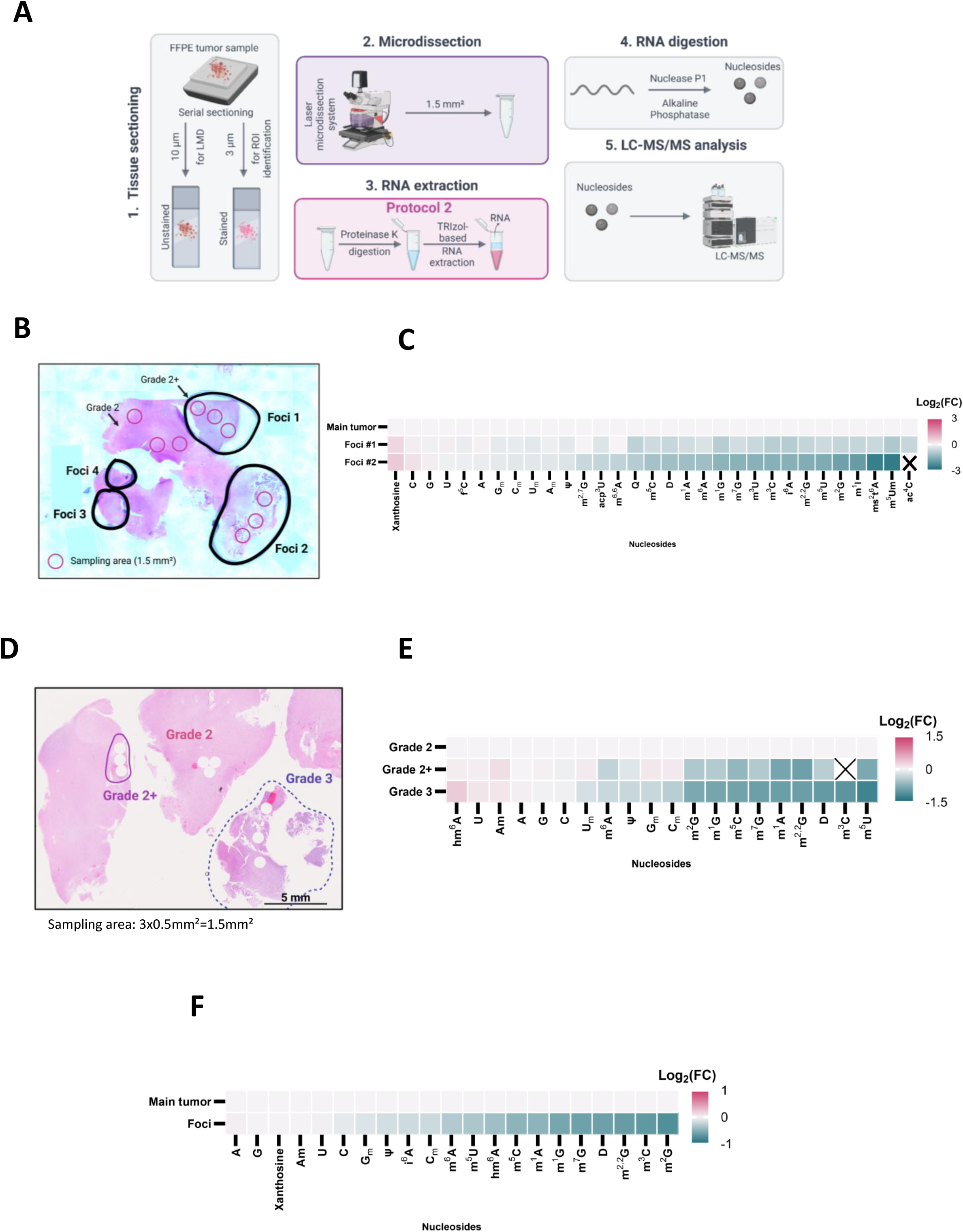
Spatial epitranscriptomic profiling of diffuse glioma reveals localized remodeling of RNA modifications during malignant progression. A. Schematic overview of the SPEAR workflow: (1) regions of interest (ROIs) were identified in FFPE diffuse glioma sections, (2) isolated by laser microdissection (LMD), (3) subjected to RNA extraction using Protocol 2, (4) enzymatically digested to individual nucleosides, and (5) analyzed by quantitative LC-MS/MS. B. Representative hematoxylin and eosin (H&E)-stained section of a WHO grade 2 oligodendroglioma (patient #2) containing focal regions of early malignant transformation (grade 2+: black outlines). Pink outlines indicate the microdissected ROIs used for LC-MS/MS analysis (original magnification, 1.25x). C. Heatmap showing relative RNA modification abundance in grade 2+ transformation foci compared with the corresponding adjacent WHO grade 2 tumor region. Values are expressed as log2 fold changes (FC) (n = 3). D. Representative H&E-stained section of a WHO grade 2 oligodendroglioma (patient #5) containing adjacent grade 2+ and WHO grade 3 foci. E. Heatmap showing relative RNA modification abundance across histological progression from WHO grade 2 to grade 2+ and WHO grade 3 (n = 3 sections). F. Heatmap showing relative RNA modification abundance in higher-grade transformation foci compared with the corresponding adjacent WHO grade 2 tumor regions across eight WHO grade 2 IDH-mutant diffuse gliomas. Values are expressed as log2 fold changes (FC).

To establish proof of concept and assess analytical reproducibility, we first applied SPEAR to a sample from patient #2, an oligodendroglioma containing four independent grade 2+ foci (**Figure 4B**). Approximately 1.6 mm² of tissue was collected from each ROI by laser microdissection, corresponding on average to 2,000-4,500 cells, depending on the tumor region sampled (**Supplementary Figure 5**). Three independent microdissections were obtained from both the primary WHO grade 2 tumor and foci 1 and 2 within the same tissue section, providing technical replicates for spatial profiling. Because of their limited size, transformation foci 3 and 4 could not accommodate repeated sampling and were therefore excluded from subsequent analyses. Despite their close spatial proximity, the primary WHO grade 2 tumor and the grade 2+ foci exhibited clearly distinct epitranscriptomic landscapes: the grade 2+ foci displayed a coordinated reduction in multiple methylated RNA modifications (**Figure 4C and Supplementary Figure 6**). In contrast, only subtle differences were detected among the individual grade 2+ foci, involving a limited number of RNA modifications, including m^6^A and m^2,2^G, suggesting more limited regional heterogeneity among these early transformation lesions.

We next asked whether SPEAR could resolve progressive epitranscriptomic remodeling across successive stages of malignant progression. A second oligodendroglioma containing spatially distinct WHO grade 2, grade 2+ and WHO grade 3 regions was therefore analyzed (patient #5) (**Figure 4D**). Three consecutive FFPE tissue sections were processed independently, and one 1.6 mm² microdissection was collected from each ROI in every section, generating three independent technical replicates for each anatomical region. Consistent with the first case, several methylated RNA modifications were reduced in the grade 2+ foci relative to the surrounding WHO grade 2 tumor, whereas these decreases became more pronounced within the WHO grade 3 foci (**Figure 4E, Supplementary Figure 7**), revealing progressive epitranscriptomic remodeling associated with focal malignant progression.

The high concordance observed among independent microdissections and serial tissue sections from these proof-of-concept experiments demonstrated the robustness and reproducibility of SPEAR. This analytical performance supported subsequent profiling of a larger validation cohort using a single representative microdissection per ROI while maximizing patient inclusion. The workflow was therefore applied to eight diffuse gliomas. Two specimens were excluded because laser microdissection failed to recover sufficient material, yielding a final cohort of six evaluable patients, including the two proof-of-concept cases. In total, 14 histologically defined ROIs, encompassing WHO grade 2 tumors together with WHO grade 3 and 4 regions, were analyzed.

Application of SPEAR across this validation cohort revealed differences in RNA-modification abundance between higher-grade foci and adjacent WHO grade 2 tissue (**Figure 4F**). Differential analysis identified multiple RNA modifications, including D, Ψ, m^5^C, m^7^G, m^5^U, and m^6^A, that showed lower abundance in higher-grade foci than in the surrounding low-grade tumor (**Supplementary Figure 8**). Despite inter-patient variability, several of these changes were observed reproducibly, across multiple independent clinical specimens, supporting the ability of SPEAR to detect localized epitranscriptomic remodeling associated with tumor progression.

## DISCUSSION

The spatial organization of the epitranscriptome remains a major unresolved challenge in RNA biology. Although RNA modifications are increasingly recognized as key regulators of RNA fate, cellular identity, plasticity, and disease progression, their investigation has largely relied on bulk analyses that average molecular signals across heterogeneous cellular populations. Conversely, sequencing-based approaches have enabled transcript-level characterization of selected RNA modifications but remain limited to individual modification classes and often require modification-specific enrichment or conversion strategies. Here, we developed SPEAR, an integrated workflow combining laser microdissection with ultrasensitive LC-MS/MS to enable quantitative spatial epitranscriptomic profiling of histologically defined FFPE tissue regions. SPEAR directly measures the nucleoside composition of RNA and enables multiplex quantification of dozens of RNA modifications within spatially resolved tissue compartments while preserving histological context.

The technological advance of SPEAR lies in the integration of two complementary approaches that have previously been applied independently. Laser microdissection provides accurate isolation of defined tissue regions while minimizing contamination from surrounding structures, enabling molecular interrogation of spatially heterogeneous biological systems (Espina et al. 2006; Kölble 2000). In parallel, LC-MS/MS has become a well-established approach for the comprehensive and quantitative analysis of RNA modifications because it directly measures modified nucleosides based on their chemical properties, without requiring amplification, antibody enrichment, or modification-specific biochemical reactions (Kellner et al. 2010; Su et al. 2014; Relier et al. 2022; Jones et al. 2023). By combining these capabilities within a workflow optimized for microscale FFPE samples, SPEAR extends epitranscriptomic analysis beyond homogenized tissues and enables direct investigation of the RNA chemical landscape within its histological context. Importantly, this integration required substantial optimization of both sample preparation and chromatographic performance to overcome the analytical constraints imposed by the minute amounts of RNA recovered from microdissected tissue.

Recent studies have begun demonstrating that RNA modifications are spatially organized within tissues, highlighting an emerging role for epitranscriptomic regulation in cellular specialization and disease (Shafik et al. 2026; Lee et al. 2022) A recent multi-region analysis of the human brain used low-input m^6^A sequencing (MeRIP-seq), combined with computational correction, orthogonal chemical validation, and genetic integration, to characterize spatial and temporal variation of m^6^A across anatomically distinct brain regions (Shafik et al. 2026). This study demonstrated that the m^6^A landscape displays strong regional organization and that spatially variable m^6^A sites are associated with genes and genetic loci implicated in neurological disorders. Similarly, Select-seq established that A-to-I RNA editing is spatially compartmentalized within cancer stem cell microniches in triple-negative breast cancer and identified relevant editing events associated with ferroptosis regulation and tumor progression (Lee et al. 2022). These studies provided important evidence that RNA modification landscapes are not uniform across tissues but instead reflect local cellular states and microenvironmental conditions. However, both approaches remain focused on individual RNA modification processes. In contrast, SPEAR provides a complementary strategy by directly quantifying the nucleoside-level composition of RNA and simultaneously profiling multiple RNA modifications within the same histological region. Future integration of SPEAR with spatial transcriptomics, spatial proteomics, metabolomics, and targeted modification-mapping approaches should enable a more comprehensive understanding of how RNA chemistry contributes to tissue organization and cellular adaptation.

Beyond its methodological contribution, SPEAR provides a spatial view of the molecular evolution of diffuse glioma. By comparing spatially distinct regions within the same tumor specimen, our approach reduces inter-patient variability and enables the identification of molecular alterations associated with focal malignant progression. Higher-grade regions displayed coordinated decreases in multiple RNA modifications, including m^5^C, m^5^U, m^6^A, hm^6^A, m^7^G, Ψ, and D. Rather than representing isolated alterations of individual RNA modification pathways, these changes are consistent with broader remodeling of the RNA chemical landscape during malignant evolution. Such spatially restricted epitranscriptomic alterations may be diluted or obscured using conventional bulk approaches, highlighting the importance of preserving tissue context when studying molecular heterogeneity. The observation of similar alterations across spatially distinct higher-grade regions and independent patient specimens supports the reproducibility of these findings, although larger cohorts will be required to determine their generality.

The coordinated reduction of several methylated RNA nucleosides during malignant transformation is particularly intriguing. Progressive global DNA hypomethylation is a well-established feature of diffuse glioma progression and is associated with acquisition of higher histological grades (Ferreyra Vega et al. 2023). Although DNA and RNA methylation are mediated by distinct molecular machineries, both processes depend on cellular methyl-donor availability, particularly S-adenosylmethionine metabolism. Our findings raise the possibility that epigenetic and epitranscriptomic remodeling may be interconnected during tumor evolution through shared metabolic constraints or coordinated regulatory mechanisms. Crosstalk between these two layers of gene expression regulation has been described in other contexts (Li et al. 2025, 2024). Alternatively, altered expression or activity of RNA-modifying enzymes may contribute independently to the observed reduction in RNA methylation. Future studies integrating spatial epigenomics, transcriptomics, proteomics, and metabolomics will be required to define the mechanisms linking DNA and RNA methylation landscapes during malignant progression. Such analyses will be particularly important to distinguish changes driven by altered RNA-modifying enzyme activity from those resulting from differences in RNA composition, cellular state, or metabolic availability of methyl donors. Importantly, because proteinase K digestion is required for efficient RNA recovery from FFPE tissues, SPEAR does not currently allow simultaneous analysis of RNA-modifying proteins, representing an intrinsic limitation of the present workflow.

A major advantage of SPEAR is its compatibility with FFPE specimens, which represent the predominant format for long-term preservation of human tissues. Unlike many current spatial approaches requiring fresh or frozen material, SPEAR enables retrospective investigation of spatial RNA modification landscapes while maintaining histological information. This feature expands the range of biological systems accessible to spatial epitranscriptomic analysis and provides opportunities to investigate how RNA chemistry contributes to tissue heterogeneity in physiological and pathological contexts. As spatial multi-omics technologies continue to evolve, SPEAR may serve as a complementary analytical layer by adding direct quantitative measurements of RNA chemical states to spatially resolved molecular profiles. Its compatibility with archived clinical material may be particularly valuable for retrospective studies in which histopathological annotation and long-term sample availability are essential.

### Limitations

Several limitations should be considered. First, although SPEAR substantially expands the capability of spatial epitranscriptomic profiling, the current workflow remains technically demanding and requires expertise in tissue preparation, pathological annotation, laser microdissection, mass spectrometry, and computational analysis. In addition, the workflow currently involves multiple manual steps, which may limit throughput and introduce technical variability. Future improvements in automation and analytical throughput will be essential for analyzing larger patient cohorts.

Second, SPEAR provides regional rather than single-cell resolution. Although laser microdissection enables enrichment of histologically defined regions, the resulting profiles represent the composite RNA chemistry of the captured tissue compartment. In addition, RNA modifications are quantified at the nucleoside level and cannot currently be assigned to specific transcripts or nucleotide positions. Therefore, SPEAR should be considered complementary to sequencing-based approaches that provide transcript-resolved information for selected RNA modifications.

Third, the molecular mechanisms responsible for the observed epitranscriptomic remodeling remain to be determined. As emphasized aboveaddressing these mechanisms would require simultaneous measurement of other variables from the same samples, such as targeted proteomics for enzyme expression and metabolomics for cofactor levels. Spatial transcriptomic and proteomic measurements could further determine whether the observed changes are associated with specific cellular populations or alterations in RNA-modifying enzyme expression.

Finally, although the spatial epitranscriptomic alterations we identified were observed consistently across multiple independent patient specimens, the cohort analyzed here remains limited in size. Validation in larger independent cohorts will therefore be necessary to establish more firmly the clinical relevance of these alterations.

## CONCLUSION

Collectively, SPEAR establishes a new technological framework for multiplexed, quantitative spatial epitranscriptomic profiling in FFPE tissues. By combining the spatial precision of laser microdissection with the chemical specificity and multiplexing capabilities of LC-MS/MS, this approach enables the quantitative interrogation of RNA modification landscapes within histologically defined tissue regions at the microscale. Applied to mouse brain and human diffuse glioma, SPEAR provided robust spatial resolution and revealed localized epitranscriptomic remodeling associated with focal malignant progression. Beyond diffuse glioma, SPEAR provides a foundation for integrating RNA chemistry into multimodal spatial biology and for dissecting how epitranscriptomic regulation contributes to tissue organization, cellular plasticity, and disease progression.

## STAR Methods

### Mouse models

Brains were collected from six 3-month-old female C57BL/6J mice. The brains were fixed for 24 h in 10% neutral-buffered formalin. Two hours after the beginning of fixation, each brain was grossly dissected into three coronal segments according to the Registry of Industrial Toxicology Animal-data (RITA) guidelines. Following fixation, the specimens were washed twice for 5 min in phosphate-buffered saline (PBS), dehydrated through ethanol, and embedded in paraffin. Dehydration and paraffin embedding were performed by RHEM. Paraffin blocks were stored at 4°C until further analysis.

### Human samples

A cohort of eight patients diagnosed with IDH-mutant, WHO grade 2 diffuse glioma (astrocytoma or oligodendroglioma) and focal high-grade transformation underwent surgical resection between 2010 and 2016 at Montpellier University Hospital (CHU de Montpellier, Montpellier, France). Archived formalin-fixed paraffin-embedded (FFPE) surgical specimens from these patients were retrieved from the Department of Pathology and Oncobiology at Montpellier University Hospital and used for this study. All tumors were reviewed and reclassified according to the 2021 World Health Organization (WHO) Classification of Tumors of the Central Nervous System. Clinical and pathological characteristics of the cohort are summarized in Table 1.

### Sectioning

Paraffin blocks were placed on a cooling plate at -10°C for at least 30 min before sectioning and maintained at this temperature until sectioning was completed. Serial sections were generated in an alternating sequence for histological staining and laser microdissection (LMD). For each paraffin block, a 10-µm-thick section designated for LMD was systematically followed by a 3-µm-thick section designated for histological staining.

The tissue sections were carefully transferred onto the surface of a 42°C water bath and allowed to expand and flatten for several seconds to minimize folding and wrinkling. The 10-µm-thick sections designated for LMD were mounted onto FrameSlides PET-Membrane (Leica, 11505190), whereas the 3-µm-thick sections designated for histological staining were mounted onto Superfrost™ Microscope Slides (Epredia, AA00008132E01MNZ20). The slides were incubated overnight at 37°C to allow complete adhesion of the tissue sections to the slides. Slides were subsequently stored in slide boxes at 4°C until further processing.

For protocol optimization experiments, 10-µm-thick sections were collected directly into 2-mL LoBind microtubes (Eppendorf, 0030 108.078) and stored at 4°C until further processing.

### Laser microdissection (LMD)

The 10-µm-thick sections mounted on FrameSlides PET-Membrane were deparaffinized by immersion in xylene for 10 min. The slides were subsequently air-dried for 5 min at room temperature under a fume hood.

The dried slides were loaded onto a Leica LMD7000 laser microdissection system, and 0.5 mL LoBind microtubes (Eppendorf, 0040 108.044) were positioned in the system’s collector holder. The caps of the collection tubes were preloaded with 25 µL of digestion buffer containing 20 mM Tris-HCl (pH 7.5), 1 mM CaCl_2_, and 0.5% SDS in ultrapure DNase/RNase free water.

ROIs were selected based on the corresponding H&E-stained sections and excised using the Leica LMD7000 at 5x magnification. The laser was operated using the following settings:

- Power: 60
- Aperture: 20
- Speed: 3
- Head current: 100%
- Pulse frequency: 3848

Microdissected material was collected directly into the digestion buffer contained in the tube caps and subsequently processed for RNA extraction.

### RNA extraction

Three RNA extraction workflows were evaluated to determine the protocol providing optimal recovery and epitranscriptomic coverage from FFPE tissue.

### Deparaffinization of sections collected directly into microtubes

For 10-µm-thick sections transferred directly into microtubes without LMD, paraffin was removed before RNA extraction. Briefly, 1 mL of xylene was added to each sample, followed by brief vortexing and incubation at 56°C for 10 min. Samples were then centrifuged at 16,900 x g for 10 min at room temperature, and the supernatant was carefully discarded. This procedure was repeated two additional times, for a total of three xylene washes, until complete deparaffinization was achieved. A final centrifugation was performed at 16,900 x g for 2 min at room temperature, and the supernatant was completely removed to eliminate residual xylene. The samples were subsequently washed with 1 mL of absolute ethanol (Carlo Erba, 4146012), vortexed, and centrifuged at 16,900 x g for 5 min at room temperature. The supernatant was discarded, followed by an additional centrifugation at 16,900 x g for 5 minutes at room temperature to remove residual ethanol. Samples were air-dried until no visible ethanol remained.

### Protocol 1: TRI Reagent-based RNA extraction

For RNA extraction using TRI Reagent, 1 mL of TRI reagent (Sigma-Aldrich, T9424) was added to the deparaffinized samples. Samples were vortexed vigorously for 2-5 s to ensure complete homogenization and incubated for 2 min at room temperature. Next, 100 µL of 1-bromo-3-chloropropane (Sigma-Aldrich, B9673) was then added, followed by thorough vortexing and incubation for 5 min at room temperature. Samples were centrifuged at 14,500 x g for 20 min at 4°C. The upper aqueous phase was carefully collected and transferred to a new 2-mL LoBind microtube. RNA was precipitated by adding 600 µL of 2-propanol (Honeywell, 33539) and 4 µL of glycogen (5 mg/mL; Invitrogen, AM9510). Samples were mixed by gentle inversion and incubated overnight at -20°C. The RNA was subsequently pelleted by centrifugation at 12,000 x g for 15 min at 4°C. The supernatant was carefully removed, and the pellet was washed with 500 µL of chilled 75% ethanol and centrifuged at 12,000 x g for 15 min at 4°C. The remaining supernatant was carefully removed without disturbing the pellet. The RNA pellet was air-dried for 5 min at 37°C and resuspended in 20 µL of DNase/RNase free water (Invitrogen, 10977-035).

### Protocol 2: Proteinase K-assisted TRI Reagent extraction

Prior to RNA extraction according to Protocol 1, samples were subjected to proteinase K pretreatment. The sample volume was adjusted to 390 µL using the digestion buffer described above (see Laser microdissection), and 10 µL of proteinase K (20 mg/mL; Invitrogen, AM2546) was added to achieve a final concentration of 500 µg/mL. Samples were vortexed briefly to ensure homogeneous mixing and incubated at 56°C for 60 min. Immediately after incubation, samples were placed on ice, transferred to a new 2-mL LoBind microtube, and processed according to Protocol 1.

### Protocol 3 : Silica column-based RNA extraction

As an alternative extraction method, total RNA was isolated using the RNeasy FFPE Kit (QIAGEN, 73504). Briefly, 240 µL of PKD buffer was added to each sample, followed by centrifugation at 11,000 x g for 1 min. Proteinase K (10 µL) was then added, and the samples were homogenized by repeated aspiration and dispensing. Samples were incubated for 15 min at 56°C, followed by 15 min at 80°C and 3 min on ice. The samples were centrifuged at full speed for 20 min at room temperature. DNase treatment was then performed by adding 25 µL of DNase Booster Buffer and 10 µL of DNase I, followed by gentle inversion and incubation for 15 min at room temperature. Next, 500 µL of RBC buffer was added and the samples were homogenized. A total of 1,120 µL of 100% ethanol was then added and mixed by repeated aspiration and dispensing. The resulting lysate was loaded onto a RNeasy MinElute spin column in successive aliquots of 700 µL. After each loading, the column was centrifuged at 8,000 x g for 15 s and the flow-through was discarded. This procedure was repeated until the entire sample had been processed. The column was then washed with 500 µL of RPE buffer and centrifuged at 8,000 x g for 30 s, followed by a second wash with 500 µL of RPE buffer and centrifugation at 8,000 x g for 2 min. The flow-through and collection tube were discarded. The column was transferred to a new 2-mL collection tube and centrifuged with the tube lid open at full speed for 5 min to remove residual wash buffer. The column was subsequently transferred to a 1.5-mL LoBind microtube (Eppendorf, 0030108051), and 22 µL of DNase/RNase-free water (Invitrogen, 10977-035) was added directly to the membrane. RNA was eluted by centrifugation at full speed for 1 min.

#### RNA quantification using Qubit

RNA concentration was determined using the Qubit RNA High-Sensitivity (HS) assay (Invitrogen, Q3285). The Qubit working solution was prepared by diluting the Qubit RNA HS reagent 1:200 in Qubit RNA HS buffer, according to the manufacturer’s instructions. For the standards, 190 µL of working solution was added to each assay tube (Invitrogen, Q32856), followed by 10 µL of the corresponding standard. For RNA samples, 195 µL of working solution was combined with 5 µL of sample. The tubes were vortexed for 5 s and incubated for 2 min at room temperature. Fluorescence was subsequently measured using a Qubit™ 4 fluorometer.

#### Reverse transcription and quantitative PCR

For reverse transcription quantitative PCR (RT-qPCR), 10 ng of total RNA was used as input for reverse transcription. Genomic DNA was removed by DNase I treatment using 1 µL of 10 X DNase buffer (Thermo Scientific, B43) and 1 µL of DNase I (Thermo Scientific, EN0521). Samples were incubated for 30 min at 37°C. Following digestion, 1 µL of 25 mM EDTA (Thermo Scientific, R1021) was added, and the samples were incubated for 10 min at 65°C and subsequently placed on ice for 1 min. For reverse transcription, 1 µL of 50 µM random hexamers (Invitrogen, 48190011) and 1 µL of 10 mM dNTP mix (Invitrogen, 10319879) were added to each sample, followed by incubation for 5 min at 65°C. Then, 4 µL of 5X First-Strand buffer, 1 µL of 0.1 M DTT, 1 µL of SuperScript III reverse transcriptase (200 U/µL; Invitrogen, 18080044) and 1 µL of RNase OUT recombinant ribonuclease inhibitor (400 U/µL; Invitrogen, 10777019) were added to each sample. Samples were then incubated at 25°C for 5 min, 50°C for 60 min, and 70°C for 15 min. The final reverse-transcription volume was 20 µL.

For qPCR, the reaction mixture was prepared using 3 µL of SYBR Green I (Roche, 04887352001), 1.9 µL of ultrapure DNase/RNase free water (Invitrogen, 10977-035) and 0.1 µL of 50 µM primer mix per well. Primers sequences are provided in Table 2. For qPCR, 1 µL of cDNA was added to each reaction. Amplification was performed using a LightCycler 480 system (Roche). Gene expression levels were normalized to 18S rRNA expression using the 2-ΔΔCt method.

**Table 2.**
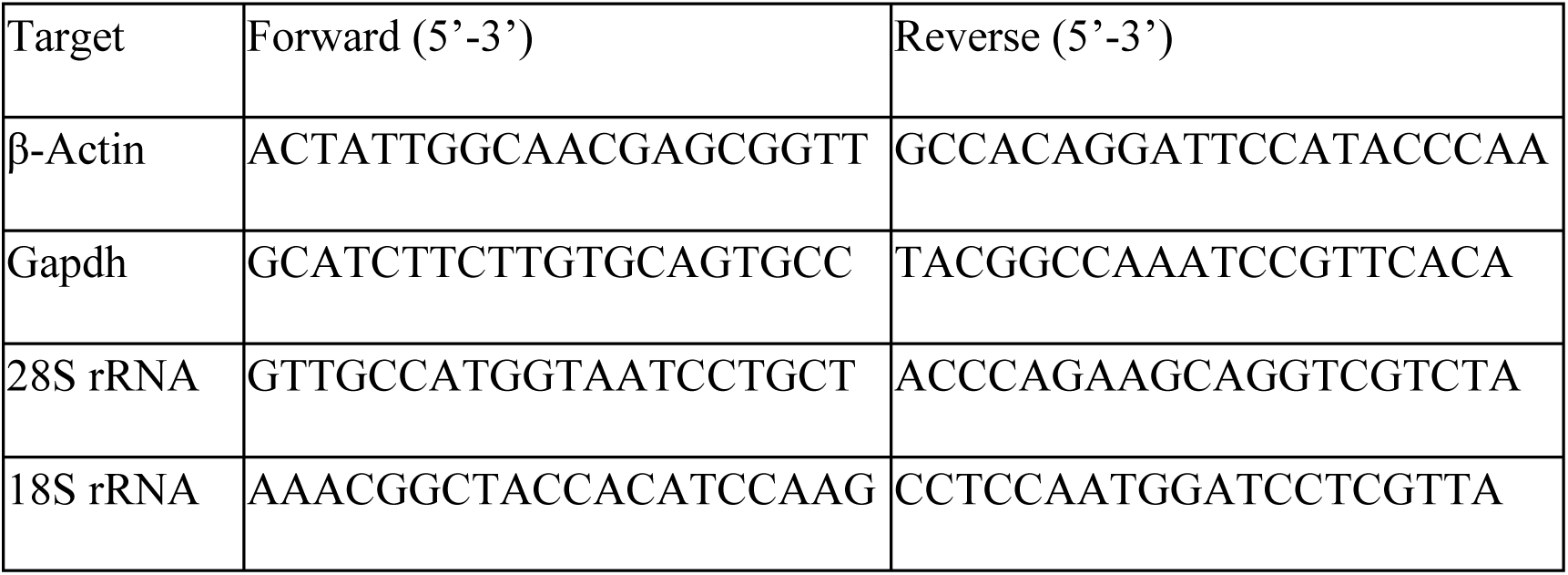
: Primer sequences.

#### RNA digestion for MS analysis

RNA samples were subjected to sequential nuclease and phosphatase digestion to generate free nucleosides for LC-MS/MS. For an initial sample volume of 20 µL, 3 µL of 100 mM ammonium acetate (NH_4_OAc), pH 5.3, and 1 µL of Nuclease P1 (1U/µL; Sigma-Aldrich, N8630) were added. Samples were incubated for 2h at 42°C. Subsequently, 3 µL of 1 M ammonium acetate and 1 µL of alkaline phosphatase from *E. coli* (1 U/µL; Sigma-Aldrich, P4252) were added, and samples were incubated for an additional 2 h at 37°C.

The resulting hydrolysates were diluted with 60 µL of mobile phase A (0.5% acetic acid) and filtered through 0.22-µm membrane filters (Millex-GV, Millipore, SLGVR04NL). A 5-µl aliquot of each hydrolysate was injected into the LC-MS/MS system.

### LC-MS/MS analysis

#### Nucleoside standards and chemicals

Nucleoside standards were purchased from Biosynth Carbosynth (Staad, Switzerland), Toronto Research Chemicals (Toronto, Canada), and Sigma-Aldrich (Steinheim, Germany). Acetonitrile (0001204101BS; CAS: 75-05-8), water (0023214102BS; CAS: 7732-18-5), ammonium acetate (001244153BS; CAS: 631-61-8) and glacial acetic acid (0001074131BS; CAS: 64-19-7) were purchased from Biosolve Chimie (Dieuze, France). All solvents and reagents used for LC-MS/MS analyses were of LC-MS grade.

#### LC Method

Nucleosides were separated using a Nexera LC system from Shimadzu. The reference method was performed on a Nexera LC-40 system equipped with a Synergi Fusion-RP column (250×2mm, 4µm, 80A, 00G-4424-B0, Phenomenex). The mobile phases consisted of 5 mM ammonium acetate pH 5.3 (mobile phase A) and acetonitrile (mobile phase B). The salt-free method was performed on a Nexera XS inert LC system equipped with a Synergi Hydro-RP column (250×2mm, 4µm, 80A, 00G-4375-B0, Phenomenex). In this configuration, ammonium acetate was replaced by 0.5% acetic acid in LC-MS-grade water as mobile phase A, while mobile phase B remained acetonitrile. For both methods, the 35-min elution gradient started at 100% mobile phase A for 3 min, followed by a linear increase in mobile phase B to 8% at 13 min and to 40% over the following 10 min. The column was then washed with 80% mobile phase B for 7 min and re-equilibrated with 100% mobile phase A for 5 min. The flow rate was 0.4 mL/min, and the oven temperature was maintained at 35°C.

### Multiple Reaction Monitoring (MRM) Method

Nucleoside detection was performed using a Shimadzu TripleQuad NX8060 mass spectrometer operated in positive electrospray ionization (ESI) mode. ESI source parameters and MRM transitions for the 46 target nucleosides are provided in **Supplementary Table 4**. The mass spectrometer was operated in dynamic MRM mode, with a 3-min retention time window and a dwell time of 20 ms.

#### Bioinformatic analysis

Due to the low RNA yield from microdissected samples, RNA concentrations could not be reliably determined before LC–MS/MS analysis, precluding normalization to RNA input. Samples in which fewer than 50% of nucleosides were detected were excluded at an initial filtering step. Among the remaining samples, we assessed both total nucleoside signal and the proportion of missing values to identify samples with potentially insufficient analytical signal. A nucleoside measurement was considered missing if it was undetected or its signal was less than twice that of the negative control. Twelve of the 13 samples with more than 10% missing values were also among the 13 samples with a total nucleoside signal below 20% of the dataset median. These 12 samples additionally separated along PC1 in principal component analysis and formed a distinct cluster in unsupervised hierarchical clustering based on Euclidean distance, consistent with a technical effect associated with low analytical signal. These observations supported the use of the proportion of missing values as a proxy for insufficient analytical signal. Accordingly, all 13 samples with more than 10% missing values were excluded before normalization.

Probabilistic quotient normalization was applied to the retained samples, followed by log2 transformation. This normalization strategy was selected because of the saturation of non-modified nucleosides in a subset of samples and a specific zone effect on U abundance between gray and white matter. Subsequent brain-wide statistical analyses were restricted to white-matter and gray-matter samples, because too few mixed-tissue samples remained in caudal and frontal FFPE blocks after quality-control filtering. When several technical sections were available, their measurements were averaged for each mouse × zone × block combination (**Supplementary Figure 3B**).

The independent contributions of anatomical block and region were evaluated using partial F-tests, comparing the complete model with reduced models excluding the term of interest. For each effect, P-values were adjusted across RNA modifications using the Benjamini–Hochberg procedure, and a false-discovery rate (FDR) below 0.05 was considered statistically significant.

To estimate the direction and magnitude of regional differences, estimated marginal means were calculated from the same models, and the adjusted contrast between gray and white matter (gray minus white matter) was reported with its standard error and 95% confidence interval **(Supplementary Table 3**). Because region comprised only two levels, this contrast tested the same hypothesis as the partial F-test for the regional effect and was not considered an additional post-hoc comparison. As the response was analyzed on the log2 scale, the estimated contrast represents a log2 fold difference.

#### Ethics statement

All animal procedures complied with European Community animal welfare guidelines (Directive 2010/63/EU). Because the animals were euthanized solely for tissue collection and no experimental procedures were performed on the animals, the study was considered non-experimental and therefore outside the scope of regulatory approval. The protocol was reviewed and approved internally by the Animal Welfare Body (SBEA) of IRCM (INSERM U1194), France.

## Acknowledgments

We thank Iria Gonzalez Dopeso-Reyes for her assistance in identifying regions of interest in murine brain sections. We acknowledge the RHEM histology facility (IRCM, Montpellier, France) for paraffin embedding of murine brains, histological staining of murine and human tissue sections, and access to the microtome used for tissue sectioning. We also thank the IRCM animal facility for animal euthanasia and murine brain collection. Finally, we acknowledge the MRI platform, particularly Sergi Navarro (CIRAD), Orestis Faklaris, and Sylvain Derossi (CRBM), for their technical advice and assistance with the laser microdissection system.

## Declaration of interests

The authors declare no competing interests.

**Supplementary Figure 1. Comparison of fold-change values obtained with the new method relative to the reference method for canonical and modified nucleosides**

Fold-change values were calculated relative to the reference method, which was set to 1. Data are presented as mean ± SD.

A. Adenosine-derived modifications.

B. Guanosine-derived modifications.

C. Cytidine-derived modifications.

The reference method used a 5 mM ammonium acetate mobile phase (pH 5.3) and a Synergi Fusion-RP column, whereas the new method used a 0.5% acetic acid mobile phase and a Synergi Hydro-RP column.

**Supplementary Figure 2: Quantification of gene expression by RT-qPCR**

A. Gene expression levels were quantified by RT-qPCR using the 2-ΔΔCt method. Data are presented as mean ± SD from three independent experiments. Statistical significance was assessed using the Mann–Whitney U test (ns, p > 0.05).

B. Representative extracted ion chromatograms illustrating the signal intensities of m^5^U, m^7^G, C_m_, and m^1^A obtained using the different RNA extraction protocols.

**Supplementary Figure 3: RNA modification profiles and sample quality control in FFPE mouse brain tissues**

A. Heatmap showing RNA modification abundance across all samples derived from FFPE mouse brain tissues. G, gray matter; W, white matter; B, mixed gray and white matter; ZM, zona moleculosa.

B. Summary of samples retained after quality control filtering.

**Supplementary Figure 4: Principal component analysis (PCA) showing the distribution of gray and white matter samples derived from FFPE mouse brain tissues**

**Supplementary Figure 5: Cell density in human diffuse glioma samples**

**Supplementary Figure 6: Spatial epitranscriptomic profiling of diffuse glioma reveals localized remodeling of RNA modifications during malignant progression.**

Nucleoside abundance was quantified in samples from the main tumor (WHO grade 2) and distinct grade 2+ transformation foci from a single patient, using LC-MS/MS in technical triplicates (n = 3). Data were normalized to the summed abundance of the canonical, unmodified nucleosides (A + C + G + U) and centered relative to the mean of the main tumor samples. Bars represent mean ± SD, with individual data points shown for each measurement. Data normality was assessed using the Shapiro–Wilk test. Statistical differences between groups were evaluated using one-way ANOVA followed by Tukey’s multiple-comparisons test for normally distributed data, or the Kruskal–Wallis test followed by Dunn’s multiple-comparisons test for non-normally distributed data (*p < 0.05; **p < 0.01; ***p < 0.001; ns, not significant).

**Supplementary Figure 7: Spatial epitranscriptomic profiling of diffuse glioma reveals localized remodeling of RNA modifications during malignant progression.**

Nucleoside abundance was quantified using LC-MS/MS and normalized to the summed abundance of the canonical, unmodified nucleosides (A, C, G, and U) in technical replicates (n = 3) from grade 2, grade 2+, and grade 3 regions of the same patient. Data were centered relative to the mean of the grade 2 samples. Bars represent mean ± SD, with individual data points shown for each measurement. Data normality was assessed using the Shapiro–Wilk test. Statistical differences between groups were evaluated using one-way ANOVA followed by Tukey’s multiple-comparisons test for normally distributed data, or the Kruskal–Wallis test followed by Dunn’s multiple-comparisons test for non-normally distributed data (*p < 0.05; **p < 0.01; ***p < 0.001; ****p < 0.0001; ns, not significant).

**Supplementary Figure 8: Comparison of relative RNA modification abundance between main tumor and malignant transformation foci**

RNA modifications in samples from the main tumor and malignant transformation foci were quantified by LC–MS/MS and normalized to the summed abundance of the canonical, unmodified nucleosides (A + C + G + U). Eight patients were initially analyzed; due to technical failures, the main tumor sample from patient #3 and the malignant transformation foci sample from patient #6 were excluded, resulting in six patients with paired samples available for comparative analysis (n = 6). Relative modification abundances were centered relative to the mean value of the main tumor samples. Lines connect paired samples originating from the same patient. Data normality was assessed using the Shapiro–Wilk test. Statistical significance between paired samples with non-normally distributed data was assessed using the two-tailed Wilcoxon signed-rank test (ns, not significant; \**p* < 0.05).

**Supplementary Table 1: Effect of RNA extraction protocol on nucleoside detection by SPEAR.**

Number of canonical and modified nucleosides detected following RNA extraction using Protocol 1, Protocol 2, or Protocol 3.

**Supplementary Table 2: Summary of samples removed from the analysis based on the quality-control threshold (detection of <90% of nucleosides per sample)**

**Supplementary Table 3: Statistical analysis of nucleoside abundance between gray and white matter**

The independent contribution of anatomical region (zone) was evaluated using partial F-tests, comparing the complete model with reduced models excluding the zone term. P-values were adjusted across RNA modifications using the Benjamini–Hochberg procedure, and a false-discovery rate below 0.05 was considered statistically significant.

To estimate the direction and magnitude of regional differences, estimated marginal means were calculated from the same models, and the adjusted contrast between gray and white matter (gray minus white matter) was reported with its standard error and 95% confidence interval. As the response was analyzed on the log2 scale, the estimated contrast represents a log2 fold difference.

**Supplementary Table 4: Mass spectrometry parameters for LC-MRM acquisition**

A. ESI instrument settings used for nucleoside analysis by LC-MS/MS.

B. Optimized multiple reaction monitoring (MRM) transitions and mass spectrometry parameters for the detection of canonical and modified nucleosides. For each analyte, the precursor ion (m/z), product ion (m/z), Q1 pre-bias voltage, collision energy (CE), and Q3 pre-bias voltage are reported. Parameters were experimentally optimized to maximize sensitivity and selectivity.

C. Retention times (min) of canonical and modified nucleosides obtained with the reference ammonium acetate-based method and the optimized salt-free method. Nucleosides are grouped according to their parent nucleoside family (modX). Retention times were determined under identical gradient conditions using either the reference chromatographic setup (LC-40/Synergi Fusion-RP/5 mM ammonium acetate) or the optimized salt-free configuration (Nexera XS inert/Synergi Hydro-RP/0.5% acetic acid).

## REFERENCES

Amalric, Amandine, Aurore Attina, Amandine Bastide, et al. 2024. « Mass Spectrometry-Based Pipeline for Identifying RNA Modifications Involved in a Functional Process: Application to Cancer Cell Adaptation ». Analytical Chemistry 96 (5): 1825-33. 10.1021/acs.analchem.3c02635.

Annesley, Thomas M. 2003. « Ion Suppression in Mass Spectrometry ». Clinical Chemistry 49 (7): 1041-44. 10.1373/49.7.1041.

Darlix, Amélie, Valérie Rigau, Julien Fraisse, Catherine Gozé, Michel Fabbro, et Hugues Duffau. 2020. « Postoperative Follow-up for Selected Diffuse Low-Grade Gliomas with WHO Grade III/IV Foci ». Article. Neurology 94 (8): 8. 10.1212/WNL.0000000000008877.

Delaunay, Sylvain, et Michaela Frye. 2019. « RNA Modifications Regulating Cell Fate in Cancer ». Nature Cell Biology 21 (5): 552-59. 10.1038/s41556-019-0319-0.

Espina, Virginia, Julia D. Wulfkuhle, Valerie S. Calvert, et al. 2006. « Laser-Capture Microdissection ». Nature Protocols 1 (2): 586-603. 10.1038/nprot.2006.85.

Ferreyra Vega, Sandra, Thomas Olsson Bontell, Teresia Kling, Asgeir Store Jakola, et Helena Carén. 2023. « Longitudinal DNA Methylation Analysis of Adult-Type IDH-Mutant Gliomas ». Acta Neuropathologica Communications 11 (1): 23. 10.1186/s40478-023-01520-1.

Guimaraes, Guilherme J., et Michael G. Bartlett. 2023. « Managing Nonspecific Adsorption to Liquid Chromatography Hardware: A Review ». Analytica Chimica Acta 1250 (avril): 340994. 10.1016/j.aca.2023.340994.

Heiss, Matthias, Felix Hagelskamp, Virginie Marchand, Yuri Motorin, et Stefanie Kellner. 2021. « Cell Culture NAIL-MS Allows Insight into Human tRNA and rRNA Modification Dynamics in Vivo ». Nature Communications 12 (1): 389. 10.1038/s41467-020-20576-4.

Jones, Joshua D., Monika K. Franco, Tyler J. Smith, et al. 2023. « Methylated guanosine and uridine modifications in S. cerevisiae mRNAs modulate translation elongation ». RSC Chemical Biology 4 (5): 363-78. 10.1039/d2cb00229a.

Jonkhout, Nicky, Julia Tran, Martin A. Smith, Nicole Schonrock, John S. Mattick, et Eva Maria Novoa. 2017. « The RNA Modification Landscape in Human Disease ». RNA 23 (12): 1754-69. 10.1261/rna.063503.117.

Kashofer, Karl, Christian Viertler, Martin Pichler, et Kurt Zatloukal. 2013. « Quality Control of RNA Preservation and Extraction from Paraffin-Embedded Tissue: Implications for RT-PCR and Microarray Analysis ». PLOS ONE 8 (7): e70714. 10.1371/journal.pone.0070714.

Kellner, Stefanie, Jürgen Burhenne, et Mark Helm. 2010. « Detection of RNA Modifications ». RNA Biology 7 (2): 237-47. 10.4161/rna.7.2.11468.

Kölble, K. 2000. « The LEICA Microdissection System: Design and Applications ». Journal of Molecular Medicine (Berlin, Germany) 78 (7): B24–25.

Lee, Amos C., Yongju Lee, Ahyoun Choi, et al. 2022. « Spatial Epitranscriptomics Reveals A-to-I Editome Specific to Cancer Stem Cell Microniches ». Nature Communications 13 (1): 2540. 10.1038/s41467-022-30299-3.

Li, Chengyu, Kexuan Chen, Qianchen Fang, et al. 2024. « Crosstalk between Epitranscriptomic and Epigenomic Modifications and Its Implication in Human Diseases ». Cell Genomics 4 (8): 100605. 10.1016/j.xgen.2024.100605.

Li, Chengyu, Kexuan Chen, Xiaoyu Li, et Xushen Xiong. 2025. « Epitranscriptome–Epigenome Interactions in Development and Disease Mechanisms ». Trends in Genetics 41 (8): 691-705. 10.1016/j.tig.2025.04.009.

Maldonado, Kenia A., et Khalid Alsayouri. 2023. « Physiology, Brain ». In StatPearls. StatPearls Publishing. http://www.ncbi.nlm.nih.gov/books/NBK551718/.

Pedeutour-Braccini, Zoé, Fanny Burel-Vandenbos, Catherine Gozé, et al. 2015. « Microfoci of Malignant Progression in Diffuse Low-Grade Gliomas: Towards the Creation of an Intermediate Grade in Glioma Classification? » Virchows Archiv 466 (4): 433-44. 10.1007/s00428-014-1712-5.

Peng, Yan, Liang Xue, Yujie Zhang, et al. 2026. « DREAMS Illuminates Spatial DNA and RNA Modification Landscapes ». Nature Cell Biology, août 24, 1-15. 10.1038/s41556-026-02038-z.

Relier, S., A. Amalric, A. Attina, et al. 2022. « Multivariate Analysis of RNA Chemistry Marks Uncovers Epitranscriptomics-Based Biomarker Signature for Adult Diffuse Glioma Diagnostics ». Analytical Chemistry 94 (35): 11967-72. 10.1021/acs.analchem.2c01526.

Shafik, Andrew M., Yong Peng, Zijie Zhang, et al. 2026. « Multi-Region m6A Epitranscriptome Profiling of the Human Brain Reveals Spatial and Temporal Variation and Enrichment of Disease-Associated Loci ». Nature Neuroscience 29 (1): 195-205. 10.1038/s41593-025-02112-z.

Sordyl, Dominik, Etienne Boileau, Agata Bernat, et al. 2026. « MODOMICS: a database of RNA modifications and related information. 2025 update and 20th anniversary ». Nucleic Acids Research 54 (D1): D219-25. 10.1093/nar/gkaf1284.

Su, Dan, Clement T. Y. Chan, Chen Gu, et al. 2014. « Quantitative Analysis of Ribonucleoside Modifications in tRNA by HPLC-Coupled Mass Spectrometry ». Nature Protocols 9 (4): 828-41. 10.1038/nprot.2014.047.

Venteicher, Andrew S., Itay Tirosh, Christine Hebert, et al. 2017. « Decoupling genetics, lineages, and microenvironment in IDH-mutant gliomas by single-cell RNA-seq ». Science (New York, N.Y.) 355 (6332): eaai8478. 10.1126/science.aai8478.

Xu, Kun, Xiang Huan, et Yu Xiao. 2026. « m6A-ARTR-DBiT for Spatial Mapping of m6A Epitranscriptome ». ACS Chemical Biology 21 (9): 2093-96. 10.1021/acschembio.6c00603.

