## Supplemental Figures and tables for "SPEAR: Spatial Epitranscriptomic Analysis by Mass Spectrometry Reveals Localized RNA Modification Remodeling During Glioma Progression"

### Supplementary Figure 1

**A**

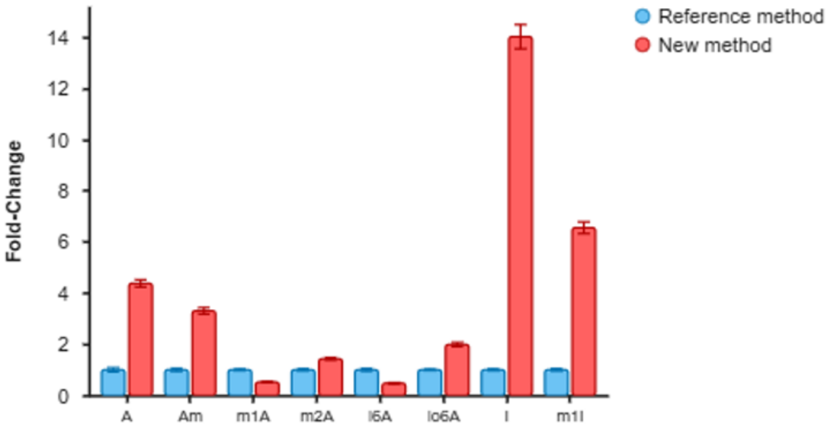


**B**

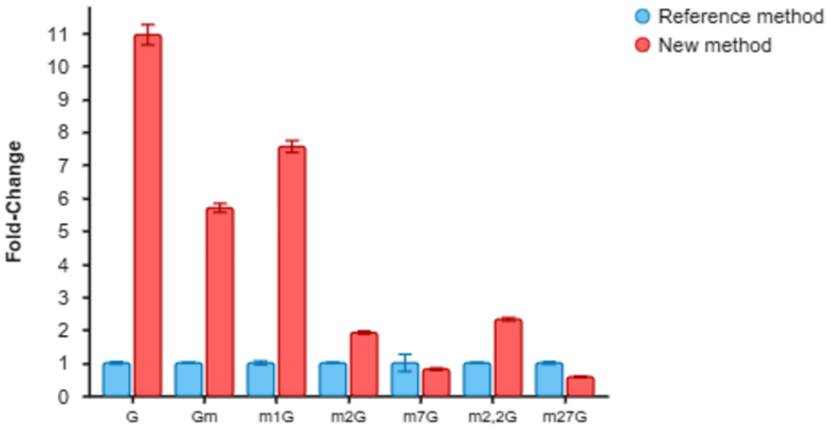


**C**

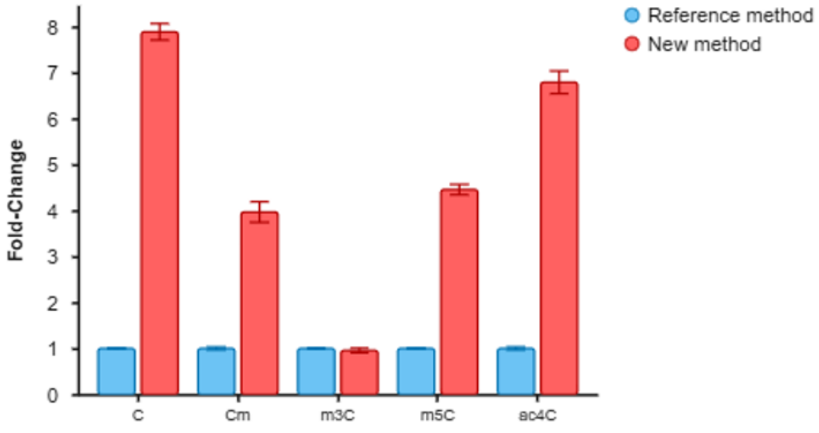

#### Supplementary Figure 2

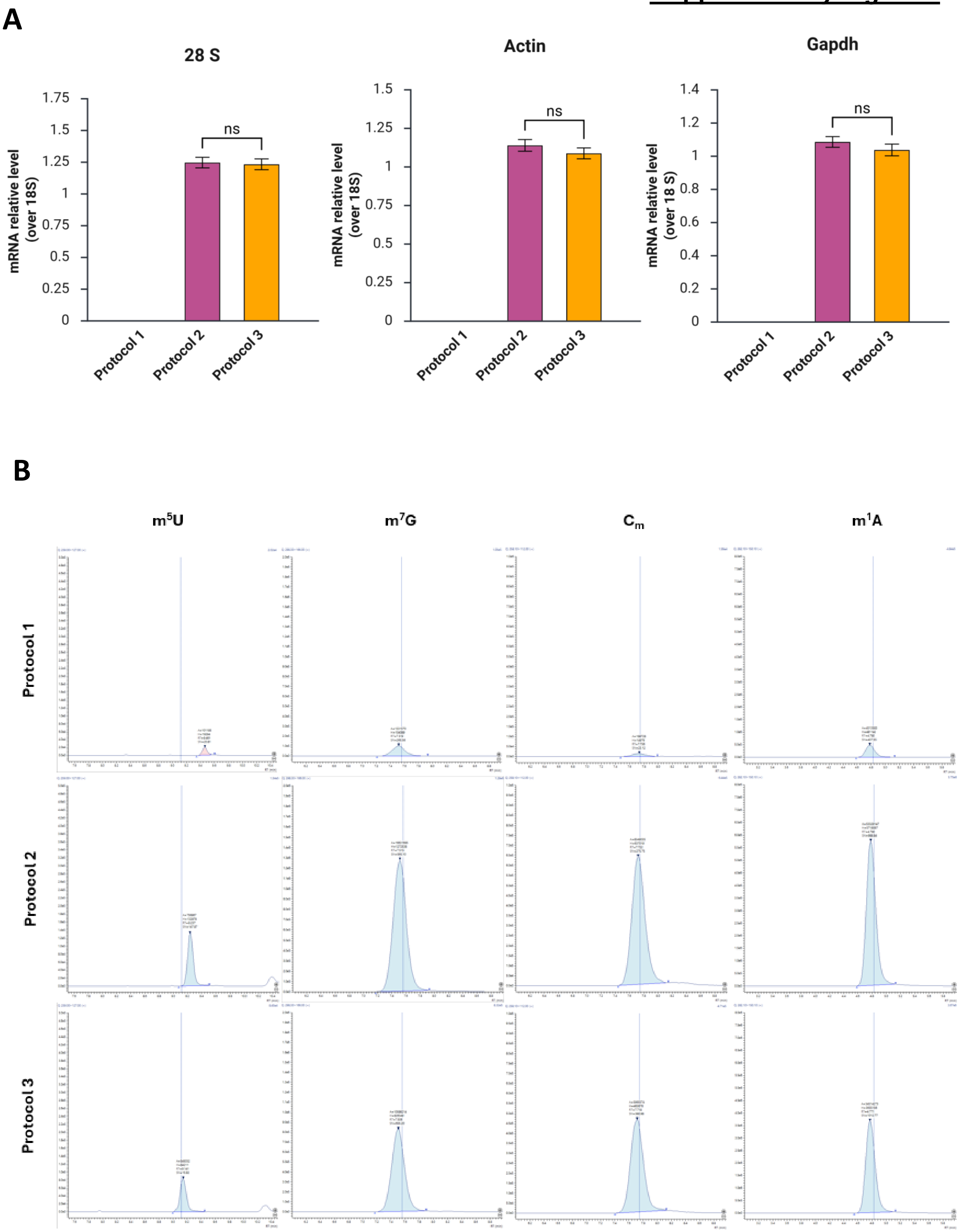

### Supplementary Figure 3

A

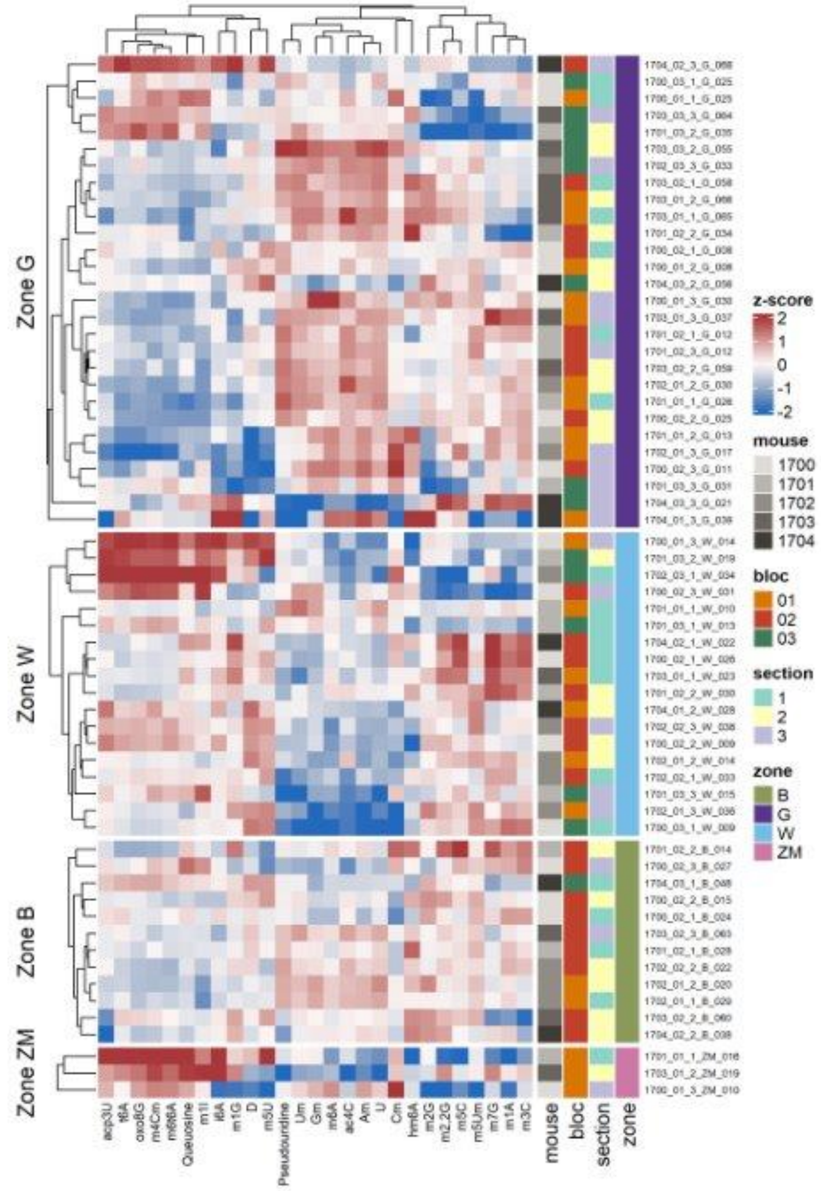


B

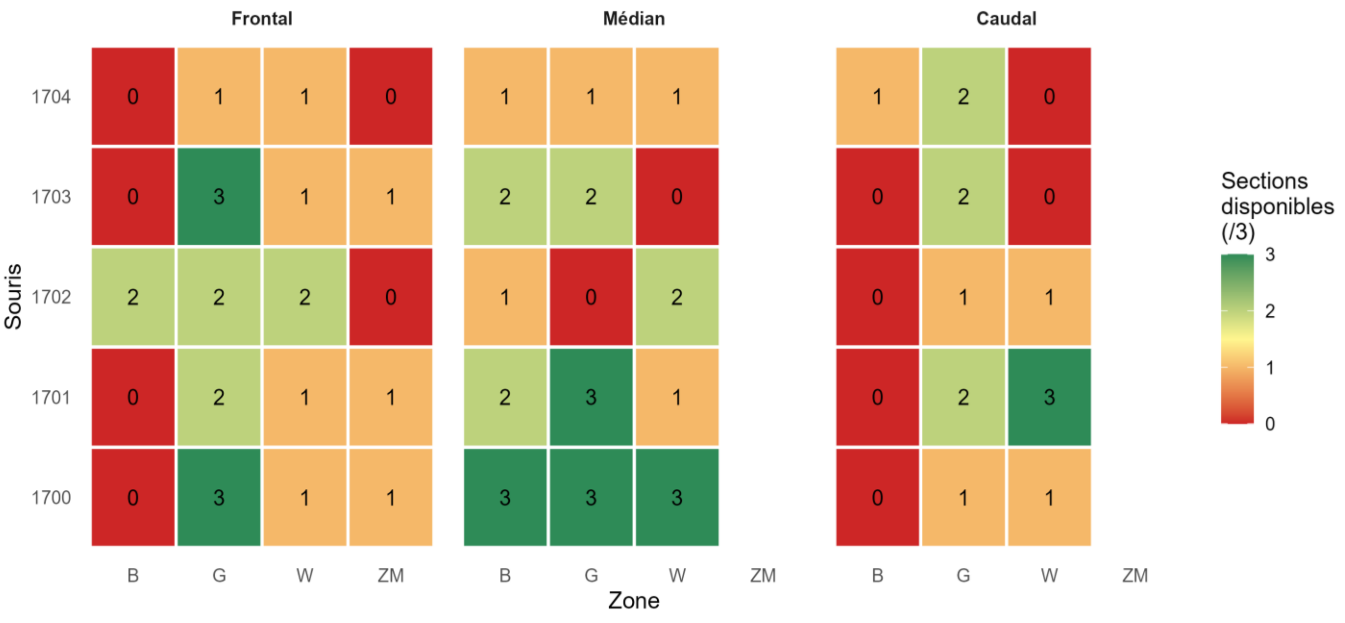

**Supplementary Figure 4**

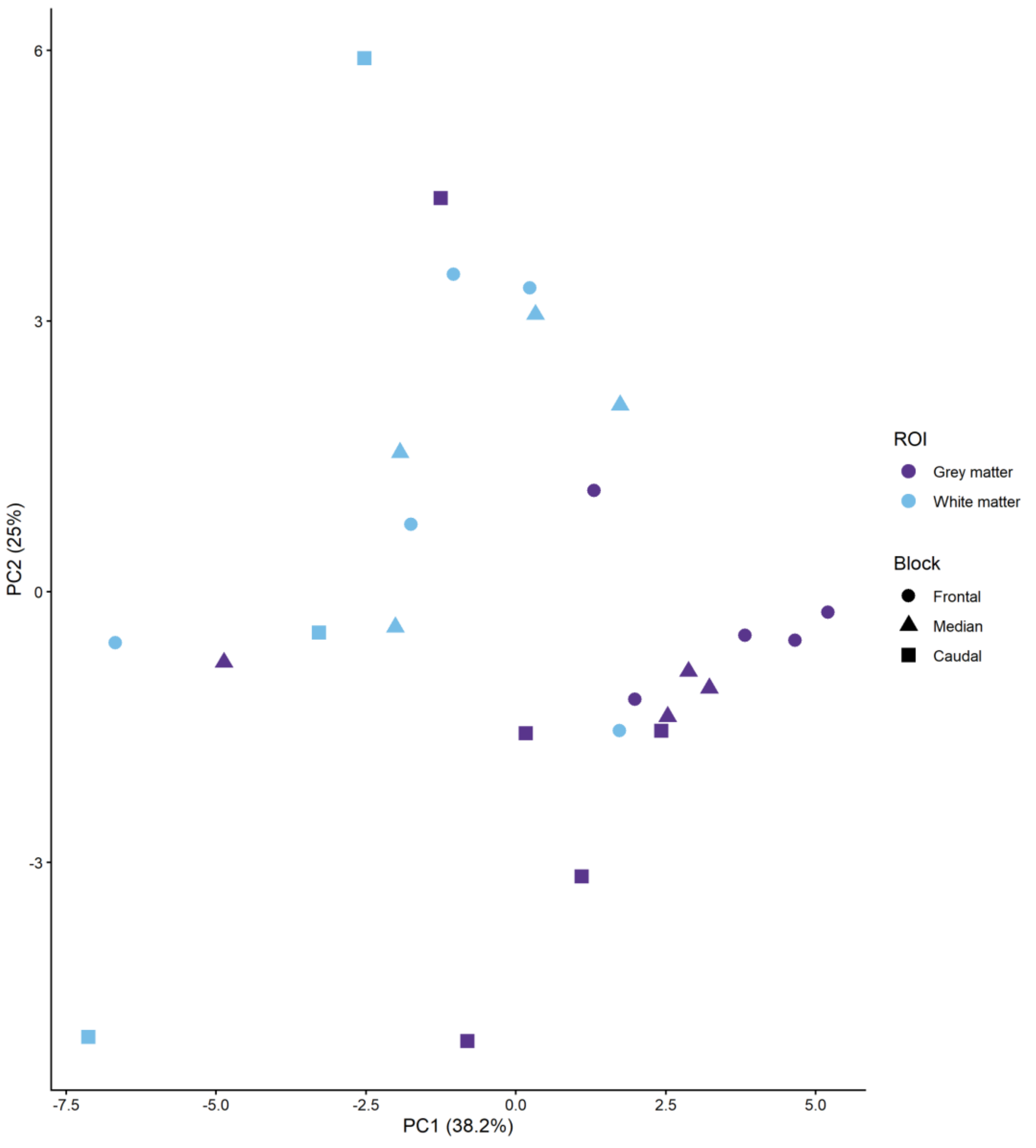

### Supplementary Figure 5

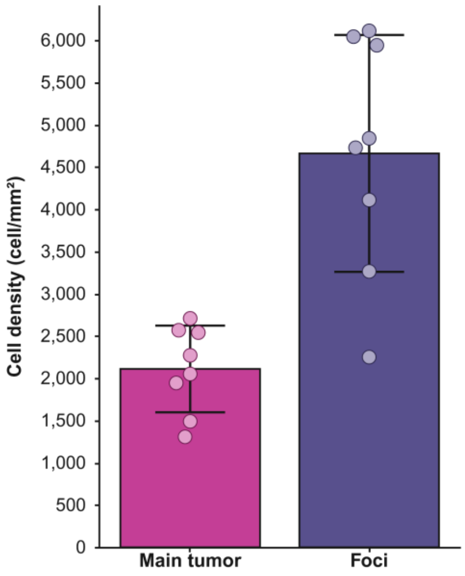


| | WHO Grade 2 main tumor | | | | Foci | | | |
|---|---|---|---|---|---|---|---|---|
| | Nb of cells | Surface (µm²) | Surface (mm²) | cells/mm² | Nb of cells | Surface (µm²) | Surface (mm²) | cells/mm² |
| Patient 1 | 520 | 170441,4761 | 0,170441476 | 3051 | 906 | 190033,5652 | 0,190033565 | 4768 |
| | 491 | 199458,948 | 0,199458948 | 2462 | 878 | 200727,8576 | 0,200727858 | 4374 |
| | 448 | 204258,9246 | 0,204258925 | 2193 | 876 | 162891,1245 | 0,162891125 | 5378 |
| Patient 2 | 362 | 169467,3364 | 0,169467336 | 2136 | 1113 | 193568,5391 | 0,193568539 | 5750 |
| | 411 | 221869,6233 | 0,221869623 | 1852 | 871 | 146100,8285 | 0,146100829 | 5962 |
| | 542 | 250466,2654 | 0,250466265 | 2164 | 1179 | 192907,7338 | 0,192907734 | 6112 |
| Patient 3 | 233 | 174116,2198 | 0,17411622 | 1338 | 410 | 181448,7122 | 0,181448712 | 2260 |
| | 185 | 169033,5227 | 0,169033523 | 1094 | 382 | 173107,504 | 0,173107504 | 2207 |
| | 309 | 207512,6009 | 0,207512601 | 1489 | 413 | 180529,8065 | 0,180529807 | 2288 |
| Patient 4 | 689 | 248683,2485 | 0,248683249 | 2771 | 591 | 175895,4275 | 0,175895428 | 3360 |
| | 456 | 167145,8005 | 0,167145801 | 2728 | 553 | 167917,5621 | 0,167917562 | 3293 |
| | 383 | 180181,7006 | 0,180181701 | 2126 | 596 | 189712,4282 | 0,189712428 | 3142 |
| Patient 5 | 355 | 185159,9367 | 0,185159937 | 1917 | 892 | 143560,4567 | 0,143560457 | 6213 |
| | 248 | 125576,1596 | 0,12557616 | 1975 | 1117 | 182400,4359 | 0,182400436 | 6124 |
| | 342 | 176066,1596 | 0,17606616 | 1942 | 862 | 143713,7053 | 0,143713705 | 5998 |
| Patient 6 | 244 | 170596,141 | 0,170596141 | 1430 | 1063 | 239918,2868 | 0,239918287 | 4431 |
| | 331 | 212113,5269 | 0,212113527 | 1560 | 853 | 186190,7266 | 0,186190727 | 4581 |
| | 355 | 239911,3521 | 0,239911352 | 1480 | 678 | 131116,4537 | 0,131116454 | 5171 |
| Patient 7 | 372 | 188958,6272 | 0,188958627 | 1969 | 824 | 155027,8293 | 0,155027829 | 5315 |
| | 530 | 182346,1298 | 0,18234613 | 2907 | 1178 | 187361,5792 | 0,187361579 | 6287 |
| | 553 | 170113,2955 | 0,170113296 | 3251 | 795 | 121813,608 | 0,121813608 | 6526 |
| Patient 8 | 315 | 169958,5818 | 0,169958582 | 1853 | 657 | 162486,9057 | 0,162486906 | 4043 |
| | 362 | 141906,8562 | 0,141906856 | 2551 | 857 | 231075,4311 | 0,231075431 | 3709 |
| | 392 | 162483,5848 | 0,162483585 | 2413 | 710 | 155047,9011 | 0,155047901 | 4579 |

| | Main tumor (cells/mm²) | Foci (cells/mm²) |
|---|---|---|
| Patient 1 | 2569 | 4840 |
| Patient 2 | 2051 | 5941 |
| Patient 3 | 1307 | 2251 |
| Patient 4 | 2541 | 3265 |
| Patient 5 | 1945 | 6112 |
| Patient 6 | 1490 | 4728 |
| Patient 7 | 2709 | 6043 |
| Patient 8 | 2272 | 4110 |
| Mean | 2111 | 4661 |
| SD | 512,8 | 1401,4 |

### Supplementary Figure 6

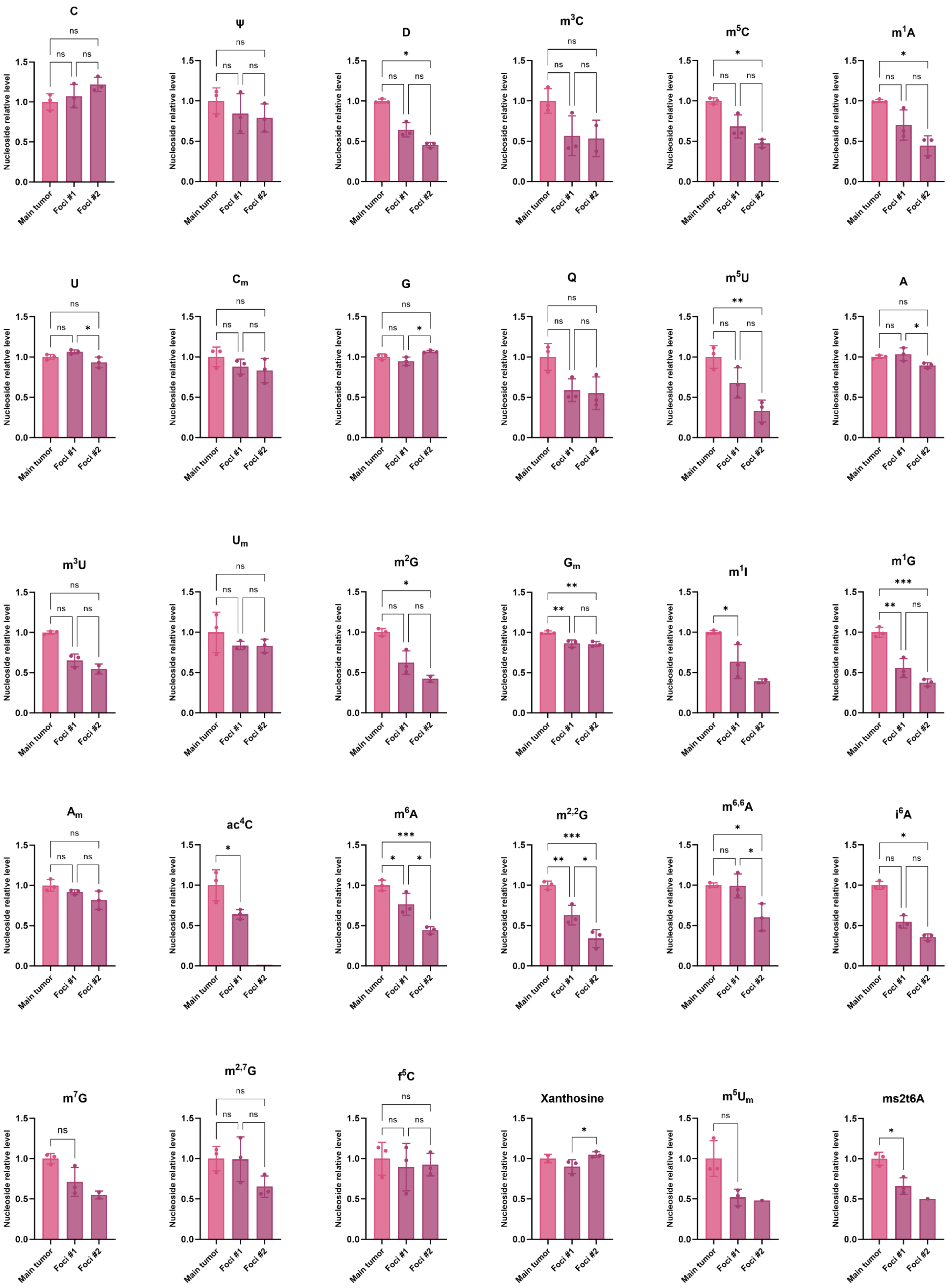

### Supplementary Figure 7

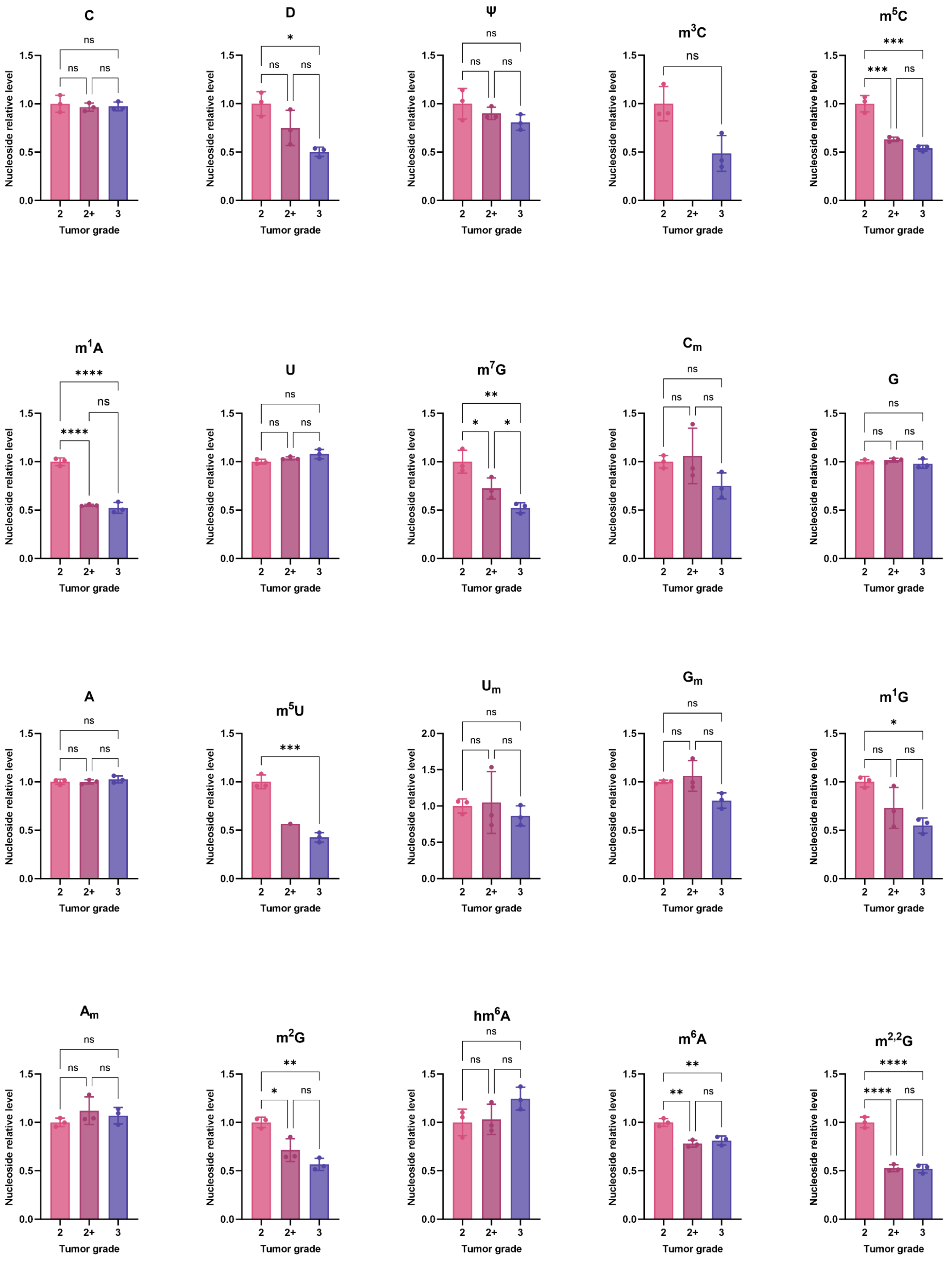

### Supplementary Figure 8

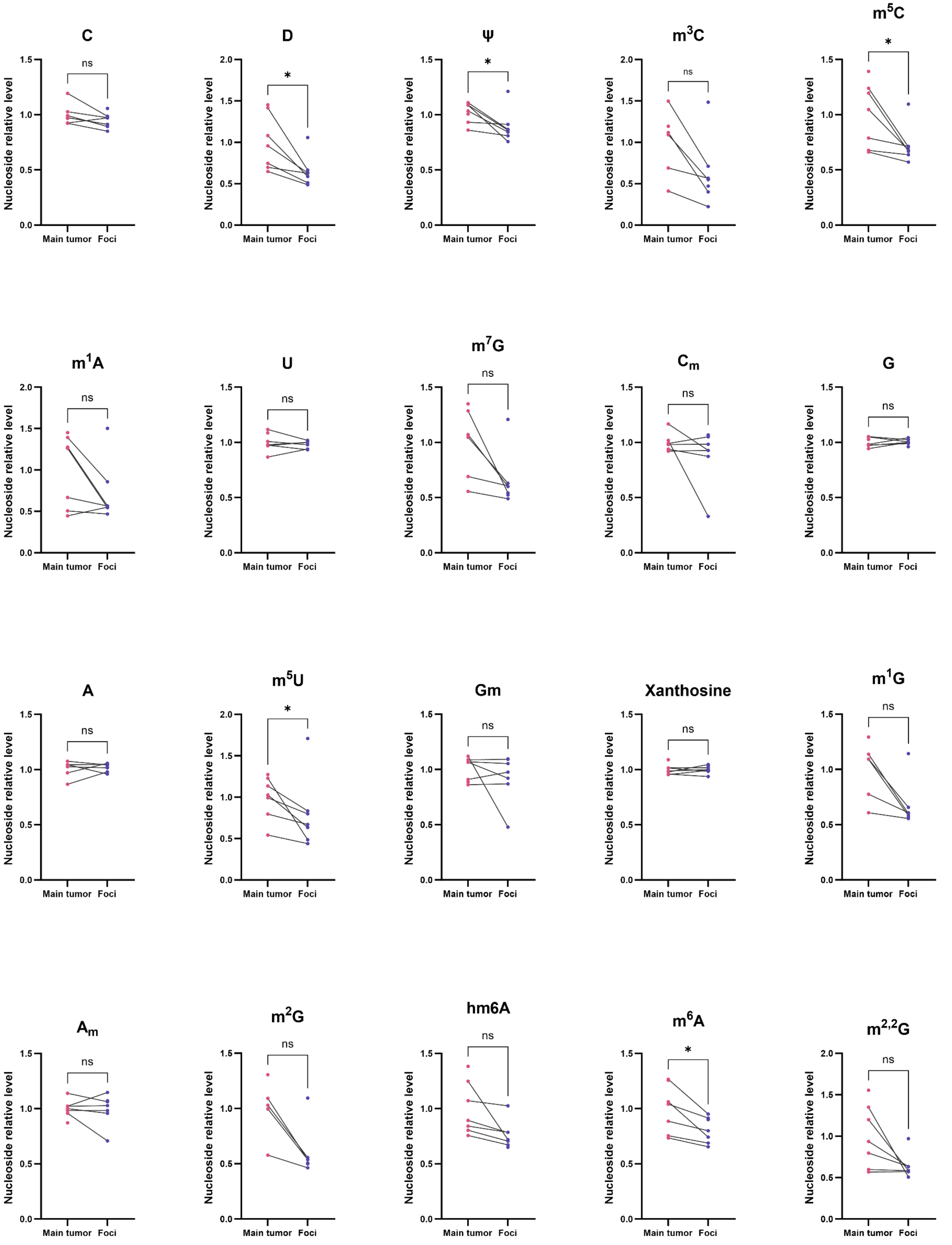

### Supplementary Table 1

| Nucleosides | Positive control (mixSTD 0.1 µM) | | | | | | Protocol 1 | | | Protocol 2 | | | Protocol 3 | | | Negative Control ($H_2O$) | Detected? | | |
|---|---|---|---|---|---|---|---|---|---|---|---|---|---|---|---|---|---|---|---|
| | 1 | 2 | 3 | 4 | 5 | 6 | n=1 | n=2 | n=3 | n=1 | n=2 | n=3 | n=1 | n=2 | n=3 | | Protocol 1 | Protocol 2 | Protocol 3 |
| C | 4956963 | 5633211 | 5348977 | 4796857 | 4936725 | 5188007 | 13085131 | 12321375 | 11384724 | 179876058 | 156432299 | 58192239 | 151667735 | 120874887 | 165377152 | 1682519 | Yes | Yes | Yes |
| D | 521709 | 429341 | 519715 | 530841 | 537208 | 486453 | 132983 | 124072 | 112568 | 1308016 | 1082195 | 205383 | 997838 | 683383 | 890778 | N/A | Yes | Yes | Yes |
| Pseudouridine | 650465 | 488527 | 589892 | 619665 | 685185 | 577505 | 229834 | 211372 | 174781 | 6503191 | 4549228 | 897368 | 4959774 | 2530576 | 6229152 | N/A | Yes | Yes | Yes |
| hm5C | 6925764 | 8218128 | 7672528 | 6957124 | 7071747 | 7531776 | N/A | N/A | N/A | 59721 | 38123 | N/A | 33921 | 17247 | 34679 | N/A | No | Yes | Yes |
| m3C | 11270689 | 13047369 | 12586507 | 11178793 | 11290422 | 12378940 | 362055 | 310243 | 275445 | 4998960 | 4099985 | 595434 | 2988512 | 1939728 | 2455633 | N/A | Yes | Yes | Yes |
| acp3U | 548698 | 586441 | 606675 | 558137 | 551412 | 600187 | 59795 | 55301 | 44856 | 690851 | 651098 | 142767 | 625823 | 317192 | 502536 | N/A | Yes | Yes | Yes |
| m1acp3Y | 1516158 | 1634529 | 1622801 | 1501186 | 1527055 | 1619530 | N/A | N/A | N/A | 41361 | 30082 | N/A | 44314 | 14232 | 85732 | N/A | No | Yes | Yes |
| ncm5U | 782172 | 737512 | 776457 | 797458 | 788053 | 765643 | 13387 | 11480 | 15416 | 205173 | 132398 | 24113 | 141338 | 64742 | 148279 | N/A | Yes | Yes | Yes |
| m1A | 14716921 | 15856137 | 16267210 | 14617751 | 14668163 | 16008493 | 4513583 | 3986694 | 3573693 | 53329147 | 38376383 | 6958712 | 34514273 | 17253584 | 32175004 | N/A | Yes | Yes | Yes |
| m5C | 2997057 | 3560814 | 3276758 | 3068195 | 2997958 | 3306199 | 1332874 | 1088941 | 1026244 | 13125873 | 8700658 | 1746659 | 8625072 | 3877376 | 9083249 | N/A | Yes | Yes | Yes |
| U | 1070750 | 988980 | 1033835 | 1022462 | 1051844 | 1030639 | 1205834 | 1119273 | 1004449 | 48690468 | 24574064 | 7343616 | 38235586 | 12183283 | 61925314 | 23196 | Yes | Yes | Yes |
| Cm | 4129453 | 4843463 | 4476380 | 3985239 | 4130878 | 4381069 | 188728 | 175999 | 163061 | 8349035 | 3728085 | 1130434 | 5965574 | 1745820 | 10307312 | 35018 | Yes | Yes | Yes |
| m7G | 9908024 | 10773392 | 10906559 | 9965507 | 10162169 | 10741431 | 1331570 | 1148035 | 1074520 | 16631695 | 10985026 | 2183030 | 10586214 | 5203198 | 10841717 | 50904 | Yes | Yes | Yes |
| S-mchm5U | 690100 | 557307 | 658627 | 761638 | 766171 | 663799 | N/A | N/A | N/A | 39540 | 20920 | N/A | 23678 | 12958 | 20323 | N/A | No | Yes | Yes |
| G | 3671947 | 4532390 | 4255730 | 3776455 | 3784245 | 4159238 | 31026188 | 27570911 | 25303257 | 226007708 | 177722155 | 131350394 | 205174451 | 139334492 | 246555041 | 1318338 | Yes | Yes | Yes |
| I | 504274 | 689788 | 645564 | 588709 | 514671 | 596531 | 1650151 | 1296903 | 1268108 | 52426991 | 26384734 | 8396502 | 40444084 | 13706715 | 65677228 | 41974 | Yes | Yes | Yes |
| A | 11614391 | 13919041 | 13193709 | 11625214 | 11634999 | 12858039 | 29454367 | 25586917 | 24249986 | 333965089 | 251380177 | 146415541 | 313273331 | 185447060 | 347664244 | 618327 | Yes | Yes | Yes |
| GalQ | 1735824 | 1885879 | 1962393 | 1713662 | 1798890 | 1980125 | 26419 | 18798 | 13215 | 176970 | 129747 | 32693 | 78833 | 53878 | 98328 | N/A | Yes | Yes | Yes |
| Queuosine | 8415449 | 9631694 | 9362467 | 8452769 | 8671005 | 9778510 | 133911 | 109288 | 121430 | 1355174 | 954864 | 212170 | 996022 | 466556 | 1035993 | N/A | Yes | Yes | Yes |
| m5U | 1056602 | 972574 | 1073619 | 1123980 | 1168274 | 1128715 | 101185 | 82553 | 67426 | 756867 | 549621 | 128100 | 545032 | 264122 | 541906 | N/A | Yes | Yes | Yes |
| Um | 534822 | 553137 | 552682 | 553961 | 544933 | 544151 | 39109 | 41563 | 33102 | 1008386 | 506421 | 129346 | 731120 | 217087 | 1189585 | N/A | Yes | Yes | Yes |
| f5C | 3277744 | 3848906 | 3634911 | 3361117 | 3369465 | 3536455 | N/A | N/A | N/A | 64783 | 32838 | 15475 | 47927 | 20334 | 82759 | N/A | No | Yes | Yes |
| m1G | 7192052 | 7967764 | 7419470 | 6858522 | 7009678 | 7285710 | 732771 | 610985 | 567339 | 8503599 | 5610313 | 1175944 | 5760780 | 2568805 | 6504288 | N/A | Yes | Yes | Yes |
| m1I | 5311055 | 5629795 | 5376247 | 5169720 | 5218842 | 5328941 | 84168 | 73722 | 67713 | 694296 | 498562 | 90364 | 417208 | 215770 | 386882 | N/A | Yes | Yes | Yes |
| m3U | 1574367 | 1616236 | 1597694 | 1575301 | 1629542 | 1610008 | N/A | N/A | N/A | 130341 | 66884 | 16159 | 89407 | 21138 | 150394 | N/A | No | Yes | Yes |
| Gm | 4897607 | 5506651 | 5330815 | 4901568 | 4946993 | 5249046 | 215894 | 238998 | 197106 | 12281393 | 5350792 | 1709119 | 8624094 | 2466912 | 17190485 | 32074 | Yes | Yes | Yes |
| Xanthosine | 1224676 | 1289154 | 1301799 | 1220567 | 1221424 | 1272651 | 2109966 | 1528421 | 1678178 | 52696931 | N/A | 9772304 | 12801235 | 14865992 | 68996872 | 72184 | Yes | Yes | Yes |
| m2G | 7358930 | 8162567 | 7812234 | 7049986 | 7146801 | 7624157 | 669277 | 598040 | 544494 | 8299136 | 5520048 | 1111614 | 5821777 | 2563067 | 6548188 | 15535 | Yes | Yes | Yes |
| Am | 7959446 | 9244686 | 8525306 | 7722739 | 8009789 | 8574398 | 157384 | 147204 | 146997 | 21472918 | 8776420 | 2868181 | 15286529 | 3953238 | 29535623 | N/A | Yes | Yes | Yes |
| ac4C | 1534495 | 1714088 | 1691698 | 1589437 | 1644287 | 1665351 | 58320 | 51643 | 49017 | 1082032 | 602626 | 140976 | 785164 | 264716 | 1074635 | N/A | Yes | Yes | Yes |
| m227G | 22199361 | 23864723 | 24031204 | 22180426 | 22737165 | 24026979 | N/A | N/A | N/A | 94108 | 74228 | 22552 | 34291 | N/A | 45849 | N/A | No | Yes | Yes |
| m6A | 8273659 | 9298461 | 8860779 | 8201415 | 8367137 | 7937021 | 49732 | 39794 | 33116 | 2549687 | 1160364 | 335851 | 1898720 | 580564 | 3323078 | N/A | Yes | Yes | Yes |
| hm6A | 731853 | 809538 | 780342 | 711897 | 712487 | 764369 | 279919 | 227657 | 230278 | 4004985 | 2491934 | 547226 | 4490518 | 1004171 | 1319961 | N/A | Yes | Yes | Yes |
| m5Um | 1089892 | 945243 | 1023076 | 1070407 | 1112056 | 1021597 | 54446 | 49082 | 40495 | 456098 | 309470 | 66945 | 296571 | 136387 | 318817 | N/A | Yes | Yes | Yes |
| m2,2G | 6992069 | 7949662 | 7699441 | 7055534 | 7099580 | 7601500 | 725421 | 621406 | 613212 | 9017473 | 5868884 | 1242245 | 5947339 | 2746044 | 6061693 | N/A | Yes | Yes | Yes |
| mcm5s2U | 1486701 | 1415830 | 1473336 | 1523588 | 1519779 | 1468072 | 17575 | 16924 | 17970 | 145046 | 102211 | 21507 | 75396 | 43733 | 72472 | N/A | Yes | Yes | Yes |
| m6Am | 8174826 | 8903051 | 8625442 | 8070334 | 8185226 | 8570819 | N/A | N/A | N/A | 143261 | 63317 | 14267 | 94002 | 17666 | 164414 | N/A | No | Yes | Yes |
| m6,6A | 15693249 | 16744163 | 16780804 | 15826045 | 15876913 | 16686776 | 54062 | 54800 | 51109 | 2959037 | 1327766 | 434420 | 2175105 | 630310 | 3835463 | 26425 | Yes | Yes | Yes |
| t6A | 5123276 | 5453137 | 5387996 | 5133436 | 5194763 | 5307814 | 306043 | 280077 | 258400 | 4012378 | 2633772 | 592706 | 2840725 | 1204611 | 2973295 | 51388 | Yes | Yes | Yes |
| m6t6A | 7906065 | 8377698 | 8226759 | 7960137 | 7968400 | 8172622 | 36215 | 31586 | 30887 | 307558 | 208749 | 48932 | 216496 | 91677 | 255097 | 9964 | Yes | Yes | Yes |
| ms2t6A | 4807999 | 5078099 | 4994236 | 4811822 | 4838195 | 4939006 | 30897 | 28100 | 25867 | 428572 | 278400 | 58351 | 253155 | 127827 | 219981 | N/A | Yes | Yes | Yes |
| i6A | 11115411 | 11894759 | 11844155 | 11157083 | 11316623 | 11774364 | 82611 | 73353 | 67508 | 1051064 | 655637 | 147034 | 722638 | 315840 | 837359 | N/A | Yes | Yes | Yes |
| ms2i6A | 184249 | 197375 | 359245 | 185927 | 181307 | 366587 | 40691 | 39691 | 39125 | 1001220 | 616607 | 136310 | 952570 | 373819 | 1120639 | 24080 | Yes | Yes | Yes |
| | | | | | | | | | | | | | | | | Number of nucleoside detected | 36 | 43 | 43 |

### Supplementary Table 2

| Sample | Action | Justification |
|---|---|---|
| 1700_01_1_G_025 | | |
| 1700_01_1_W_009 | remove | Less than 90% of nucleotides detected |
| 1700_01_1_ZM_007 | remove | Less than 90% of nucleotides detected |
| 1700_01_2_G_008 | | |
| 1700_01_2_W_023 | remove | Less than 90% of nucleotides detected |
| 1700_01_2_ZM_024 | remove | Less than 90% of nucleotides detected |
| 1700_01_3_G_030 | | |
| 1700_01_3_W_014 | | |
| 1700_01_3_ZM_010 | | |
| 1700_02_1_B_024 | | |
| 1700_02_1_G_008 | | |
| 1700_02_1_W_026 | | |
| 1700_02_2_B_015 | | |
| 1700_02_2_G_025 | | |
| 1700_02_2_W_009 | | |
| 1700_02_3_B_027 | | |
| 1700_02_3_G_011 | | |
| 1700_02_3_W_031 | | |
| 1700_03_1_G_025 | | |
| 1700_03_1_W_009 | | |
| 1701_01_1_G_026 | | |
| 1701_01_1_W_010 | | |
| 1701_01_1_ZM_016 | | |
| 1701_01_1_ZM_038 | remove | Less than 90% of nucleotides detected |
| 1701_01_2_G_013 | | |
| 1701_01_2_W_032 | remove | Less than 90% of nucleotides detected |
| 1701_01_2_ZM_029 | remove | Less than 90% of nucleotides detected |
| 1701_01_3_W_011 | remove | Less than 90% of nucleotides detected |
| 1701_02_1_B_028 | | |
| 1701_02_1_G_012 | | |
| 1701_02_1_W_027 | remove | Less than 90% of nucleotides detected |
| 1701_02_2_B_014 | | |
| 1701_02_2_G_034 | | |
| 1701_02_2_W_018 | remove | Less than 90% of nucleotides detected |
| 1701_02_2_W_030 | | |
| 1701_02_3_B_028 | remove | Less than 90% of nucleotides detected |
| 1701_02_3_G_012 | | |
| 1701_03_1_G_029 | remove | Less than 90% of nucleotides detected |
| 1701_03_1_W_013 | | |
| 1701_03_2_G_035 | | |
| 1701_03_2_W_019 | | |
| 1701_03_3_G_031 | | |
| 1701_03_3_W_015 | | |
| 1702_01_1_B_029 | | |
| 1702_01_1_G_013 | remove | Less than 90% of nucleotides detected |
| 1702_01_1_W_037 | remove | Less than 90% of nucleotides detected |
| 1702_01_2_B_020 | | |
| 1702_01_2_G_030 | | |
| 1702_01_2_W_014 | | |
| 1702_01_3_B_050 | remove | Less than 90% of nucleotides detected |
| 1702_01_3_G_017 | | |
| 1702_01_3_W_036 | | |
| 1702_02_1_B_031 | remove | Less than 90% of nucleotides detected |
| 1702_02_1_G_015 | remove | Less than 90% of nucleotides detected |
| 1702_02_1_W_033 | | |
| 1702_02_2_B_022 | | |
| 1702_02_2_G_032 | remove | Less than 90% of nucleotides detected |
| 1702_02_2_W_016 | remove | Less than 90% of nucleotides detected |
| 1702_02_3_B_039 | remove | Less than 90% of nucleotides detected |
| 1702_02_3_G_018 | remove | Less than 90% of nucleotides detected |
| 1702_02_3_W_038 | | |
| 1702_03_1_G_016 | remove | Less than 90% of nucleotides detected |
| 1702_03_1_W_034 | | |
| 1702_03_2_G_032 | remove | Less than 90% of nucleotides detected |
| 1702_03_2_W_027 | remove | Less than 90% of nucleotides detected |
| 1702_03_3_G_033 | | |
| 1702_03_3_W_017 | remove | Less than 90% of nucleotides detected |
| 1703_01_1_G_065 | | |
| 1703_01_1_W_023 | | |
| 1703_01_1_ZM_020 | remove | Less than 90% of nucleotides detected |
| 1703_01_2_G_066 | | |
| 1703_01_2_W_021 | remove | Less than 90% of nucleotides detected |
| 1703_01_2_ZM_019 | | |
| 1703_01_3_G_037 | | |
| 1703_01_3_W_034 | remove | Less than 90% of nucleotides detected |
| 1703_02_1_B_019 | remove | Less than 90% of nucleotides detected |
| 1703_02_1_G_058 | | |
| 1703_02_2_B_060 | | |
| 1703_02_2_G_059 | | |
| 1703_02_2_W_022 | remove | Less than 90% of nucleotides detected |
| 1703_02_3_B_051 | remove | Less than 90% of nucleotides detected |
| 1703_02_3_B_063 | | |
| 1703_03_1_G_035 | remove | Less than 90% of nucleotides detected |
| 1703_03_1_W_037 | remove | Less than 90% of nucleotides detected |
| 1703_03_2_G_055 | | |
| 1703_03_2_W_044 | remove | Less than 90% of nucleotides detected |
| 1703_03_3_G_064 | | |
| 1703_03_3_W_017 | remove | Less than 90% of nucleotides detected |
| 1704_01_1_B_070 | remove | Less than 90% of nucleotides detected |
| 1704_01_1_G_036 | remove | Less than 90% of nucleotides detected |
| 1704_01_1_W_020 | remove | Less than 90% of nucleotides detected |
| 1704_01_2_B_033 | remove | Less than 90% of nucleotides detected |
| 1704_01_2_G_036 | remove | Less than 90% of nucleotides detected |
| 1704_01_2_W_028 | | |
| 1704_01_3_B_040 | remove | Less than 90% of nucleotides detected |
| 1704_01_3_G_039 | | |
| 1704_01_3_W_021 | remove | Less than 90% of nucleotides detected |
| 1704_02_1_B_045 | remove | Less than 90% of nucleotides detected |
| 1704_02_1_G_018 | remove | Less than 90% of nucleotides detected |
| 1704_02_1_W_022 | | |
| 1704_02_2_B_038 | | |
| 1704_02_2_G_035 | remove | Less than 90% of nucleotides detected |
| 1704_02_2_W_024 | remove | Less than 90% of nucleotides detected |
| 1704_02_3_B_053 | remove | Less than 90% of nucleotides detected |
| 1704_02_3_G_068 | | |
| 1704_02_3_W_054 | remove | Less than 90% of nucleotides detected |
| 1704_03_1_B_048 | | |
| 1704_03_1_G_061 | remove | Less than 90% of nucleotides detected |
| 1704_03_1_W_069 | remove | Less than 90% of nucleotides detected |
| 1704_03_2_B_023 | remove | Less than 90% of nucleotides detected |
| 1704_03_2_G_056 | | |
| 1704_03_2_W_049 | remove | |
| 1704_03_3_B_026 | remove | Less than 90% of nucleotides detected |
| 1704_03_3_G_021 | | |
| 1704_03_3_W_043 | remove | Less than 90% of nucleotides detected |

### Supplementary Table 3

| Nucleosides | difference_G_W | Standard_error | IC95_inf | IC95_sup | t | p_zone | padj_zone |
|---|---|---|---|---|---|---|---|
| Am | 0,858684403 | 0,192616078 | 0,454013 | 1,263356 | 4,45801 | 0,000304 | 0,002735 |
| Cm | 0,200668444 | 0,17279059 | -0,16235 | 0,563688 | 1,161339 | 0,260674 | 0,319918 |
| D | -0,277518552 | 0,102962714 | -0,49384 | -0,0612 | -2,69533 | 0,014798 | 0,057077 |
| Gm | 0,538981055 | 0,160425574 | 0,201939 | 0,876023 | 3,359695 | 0,003489 | 0,016606 |
| Pseudouridine | 0,263547157 | 0,113177077 | 0,025771 | 0,501323 | 2,328627 | 0,031731 | 0,085675 |
| Queuosine | -0,384491469 | 0,115311884 | -0,62675 | -0,14223 | -3,33436 | 0,00369 | 0,016606 |
| U | 0,73495411 | 0,132476095 | 0,456632 | 1,013276 | 5,547824 | 2,88E-05 | 0,000389 |
| Um | 0,390241933 | 0,175190852 | 0,02218 | 0,758304 | 2,227525 | 0,03891 | 0,095506 |
| ac4C | 0,395820195 | 0,068256155 | 0,252419 | 0,539221 | 5,79904 | 1,71E-05 | 0,000389 |
| acp3U | -0,666906106 | 0,259670234 | -1,21245 | -0,12136 | -2,56828 | 0,019345 | 0,06375 |
| hm6A | 0,435556419 | 0,221706036 | -0,03023 | 0,901344 | 1,964567 | 0,065094 | 0,120671 |
| i6A | -0,125951721 | 0,073062188 | -0,27945 | 0,027546 | -1,7239 | 0,101859 | 0,160804 |
| m1A | -0,116930506 | 0,11924051 | -0,36745 | 0,133585 | -0,98063 | 0,339774 | 0,382246 |
| m1G | -0,084130076 | 0,060818985 | -0,21191 | 0,043646 | -1,38329 | 0,183495 | 0,247719 |
| m1I | -0,443650406 | 0,175818532 | -0,81303 | -0,07427 | -2,52334 | 0,02125 | 0,06375 |
| m2,2G | -0,053441774 | 0,084988168 | -0,232 | 0,125112 | -0,62881 | 0,537369 | 0,580359 |
| m2G | 0,000419697 | 0,087420753 | -0,18324 | 0,184084 | 0,004801 | 0,996222 | 0,996222 |
| m3C | -0,271671023 | 0,224403455 | -0,74313 | 0,199783 | -1,21064 | 0,241692 | 0,310747 |
| m4Cm | -0,661927152 | 0,333853191 | -1,36333 | 0,039472 | -1,98269 | 0,062876 | 0,120671 |
| m5C | -0,037861595 | 0,095993762 | -0,23954 | 0,163814 | -0,39442 | 0,697907 | 0,72475 |
| m5U | -0,176190041 | 0,114305616 | -0,41634 | 0,063957 | -1,54139 | 0,140617 | 0,199825 |
| m5Um | -0,243748302 | 0,235935862 | -0,73943 | 0,251935 | -1,03311 | 0,315233 | 0,370056 |
| m6A | 0,784474168 | 0,181769285 | 0,402591 | 1,166357 | 4,315769 | 0,000416 | 0,00281 |
| m6t6A | -0,512232638 | 0,262802222 | -1,06436 | 0,039894 | -1,94912 | 0,06704 | 0,120671 |
| m7G | -0,180004367 | 0,086041716 | -0,36077 | 0,000763 | -2,09206 | 0,050875 | 0,114469 |
| oxo8G | -0,511693498 | 0,30109699 | -1,14427 | 0,120888 | -1,69943 | 0,106454 | 0,160804 |
| t6A | -0,255134233 | 0,150474665 | -0,57127 | 0,061001 | -1,69553 | 0,107203 | 0,160804 |

### Supplementary Table 4

A

| Parameters | Setting |
|---|---|
| Nebulizing Gas Flow | 3 L/min |
| Heating Gas Flow | 10 L/min |
| Interface Temperature | 350 °C |
| DL Temperature | 225°C |
| Heat Block Temperature | 400°C |
| Drying gas flow | 3 L/min |
| Focus Voltage | 2 kV |
| Interface Voltage | 0.5 kV |
| CID Gas | 250 kPa |

B

| Name | Precursor m/z | Products m/z | Target Q1 Pre Bias (V) | Target Collision Energy (V) | Target Q3 Pre Bias (V) |
|---|---|---|---|---|---|
| A | 268 | 136 | -11 | -16 | -11 |
| ac4C | 286.1 | 154 | -12 | -11 | -13 |
| | 286.1 | 112 | -11 | -27 | -17 |
| acp3U | 346 | 214 | -14 | -17 | -12 |
| | 346 | 197 | -13 | -23 | -18 |
| | 346 | 168 | -16 | -20 | -30 |
| Am | 282 | 136 | -11 | -15 | -11 |
| C | 244.1 | 112 | -12 | -14 | -18 |
| Cm | 258.1 | 112 | -11 | -13 | -17 |
| cm5U | 303 | 171 | -12 | -10 | -14 |
| | 303 | 125 | -12 | -26 | -20 |
| | 303 | 153 | -26 | -16 | -27 |
| D | 247.1 | 115 | -19 | -10 | -18 |
| | 247.1 | 115 | -19 | -10 | -18 |
| | 247.1 | 97 | -21 | -21 | -28 |
| f5C | 272.1 | 140.1 | -13 | -12 | -12 |
| | 272.1 | 97 | -13 | -33 | -15 |
| | 272.1 | 69 | -11 | -42 | -25 |
| G | 284.1 | 152 | -14 | -14 | -13 |
| GalQ | 572 | 120.9 | -22 | -72 | -14 |
| | 572 | 120.9 | -22 | -72 | -14 |
| Gm | 298.1 | 152 | -12 | -13 | -13 |
| hm5C | 274.1 | 142 | -20 | -15 | -21 |
| hm6A | 298.1 | 136 | -14 | -20 | -11 |
| | 298.1 | 136 | -14 | -20 | -11 |
| I | 269.1 | 137 | -11 | -13 | -11 |
| i6A | 336.15 | 204 | -10 | -18 | -11 |
| | 336.15 | 148 | -17 | -25 | -23 |
| | 336.15 | 136 | -27 | -30 | -22 |
| m1A | 282.1 | 150.1 | -12 | -20 | -13 |
| m1acp3Y | 360.1 | 270 | -30 | -16 | -24 |
| | 360.1 | 139 | -17 | -32 | -21 |
| | 360.1 | 270 | -30 | -16 | -24 |
| m1G | 298 | 166 | -12 | -17 | -14 |
| m1I | 283.1 | 151 | -10 | -13 | -25 |
| m2,2G | 312 | 180 | -12 | -16 | -15 |
| m2,2,7G | 326.15 | 194 | -13 | -17 | -17 |
| m2G | 298.1 | 166 | -14 | -16 | -14 |
| m3C | 258.1 | 126 | -20 | -25 | -20 |
| m3U | 259 | 127 | -21 | -13 | -20 |
| m5C | 258 | 126 | -10 | -13 | -10 |
| m5Cm | 272.1 | 126.2 | -11 | -14 | -10 |
| m5U | 259 | 127 | -22 | -11 | -10 |
| m5Um | 273.1 | 127 | -11 | -12 | -10 |
| m6,6A | 296 | 164.1 | -12 | -20 | -14 |
| m6A | 282.1 | 150.1 | -11 | -20 | -13 |
| m6Am | 296 | 150 | -12 | -18 | -24 |
| m6t6A | 427.2 | 295 | -12 | -14 | -17 |
| m7G | 298.3 | 166 | -10 | -15 | -26 |
| mcm5s2U | 333.1 | 201 | -10 | -11 | -11 |
| | 333.1 | 169 | -16 | -19 | -15 |
| | 333.1 | 141 | -13 | -28 | -22 |
| mcm5U | 317.1 | 185 | -13 | -11 | -10 |
| | 317.1 | 153 | -15 | -18 | -25 |
| | 317.1 | 125 | -28 | -28 | -20 |
| mcm5Um | 331 | 185 | -13 | -11 | -10 |
| | 331 | 125 | -13 | -28 | -19 |
| | 331 | 153 | -27 | -17 | -25 |
| ms2t6A | 458.9 | 327.1 | -11 | -14 | -13 |
| | 458.9 | 182 | -11 | -33 | -16 |
| | 458.9 | 208 | -11 | -28 | -12 |
| | 458.9 | 134 | -11 | -55 | -21 |
| ncm5U | 302.1 | 170 | -12 | -11 | -14 |
| | 302.1 | 153 | -14 | -18 | -13 |
| | 302.1 | 125 | -25 | -28 | -19 |
| Pseudouridine | 245.05 | 209 | -10 | -10 | -12 |
| | 245.05 | 179 | -20 | -11 | -15 |
| | 245.05 | 155 | -26 | -13 | -25 |
| Queuosine | 410.2 | 163 | -10 | -24 | -26 |
| | 410.2 | 163 | -10 | -24 | -26 |
| S-mchm5U | 333 | 183 | -13 | -16 | -10 |
| | 333 | 201 | -27 | -10 | -18 |
| t6A | 413.1 | 281 | -17 | -14 | -17 |
| | 413.1 | 136 | -10 | -31 | -11 |
| | 413.1 | 162 | -10 | -27 | -26 |
| U | 245.1 | 113 | -10 | -11 | -19 |
| Um | 259.1 | 113 | -10 | -11 | -18 |
| Xanthosine | 285.2 | 153.1 | -30 | -11 | -27 |
| | 285.2 | 136 | -10 | -28 | -22 |
| | 285.2 | 153.1 | -30 | -11 | -27 |

C

| Nucleosides | Reference method | New method |
|---|---|---|
| D | 3.4 | 2.6 |
| Pseudouridine | 3.5 | 2.7 |
| m3C | 3.6 | 3.8 |
| m1A | 4.1 | 4.8 |
| PreQ1 | 4.2 | 4.9 |
| C | 4.6 | 2.7 |
| hm5C | 5.3 | 2.8 |
| ncm5U | 5.7 | 4.1 |
| m7G | 6.5 | 7.4 |
| U | 6.7 | 4.8 |
| Queuine | 8.4 | 8.7 |
| m5C | 8.5 | 4.5 |
| m5U | 8.5 | 8.8 |
| Queuosine | 8.9 | 9.2 |
| Cm | 9.1 | 7.4 |
| I | 9.3 | 8.6 |
| G | 9.6 | 8.8 |
| m27G | 9.7 | 10.0 |
| m3U | 10.0 | 9.9 |
| mo5U | 10.0 | 9.0 |
| oxo8G | 10.0 | 9.2 |
| Um | 10.8 | 9.6 |
| m1I | 11.4 | 10.6 |
| m227G | 11.4 | 11.7 |
| m2G | 11.6 | 11.1 |
| Gm | 11.8 | 10.8 |
| mcm5U | 12.1 | 10.5 |
| m1G | 12.2 | 10.5 |
| ac4C | 12.4 | 11.2 |
| A | 12.8 | 9.0 |
| ho8A | 13.0 | 11.1 |
| m22G | 14.2 | 13.1 |
| Am | 15.4 | 11.4 |
| mcm5s2U | 16.0 | 14.0 |
| m2A | 16.1 | 9.9 |
| m3Um | 16.1 | 14.6 |
| m6A | 16.5 | 12.1 |
| m6Am | 17.8 | 15.2 |
| io6A | 18.1 | 17.0 |
| m66A | 18.5 | 16.0 |
| m66Am | 19.9 | 18.0 |
| i6A | 22.1 | 20.8 |
